# The ReAct project: Analysis of data from 23 different laboratories to characterise DNA recovery given two sets of activity level propositions

**DOI:** 10.1101/2024.10.10.617148

**Authors:** Peter Gill, Ane Elida Fonneløp, Tacha Hicks, Stavroulla Xenophontos, Marios Cariolou, Roland van Oorschot, Iris Buckel, Viktorija Sukser, Sunčica Papić, Siniša Merkaš, Ana Kostic, Angela Marques Pereira, Christina Teutsch, Christina Forsberg, Cordula Haas, Elizabet Petkovski, Fabian Hass, Jan Masek, Jelena Stosic, Yong Sheng Lee, Christopher Kiu-Choong Syn, Linda Groombridge, Marc Trimborn, Marilena Hadjivassiliou, Michelle Breathnach, Jana Novackova, Walther Parson, Petra Hatzer-Grubwieser, Sanna Pietikäinen, Simone Joas, Sascha Willuweit, Stefanie Grethe, Tamara Milićević, Therese Hasselqvist, Venus Kallupurackal, Vlastimil Stenzl, Staffan Jansson, Ingrun Glocker, Sarah Brunck, Karoline Nyhagen, Anne Berit Dyve Lingelem, Heli Autere, Devon Thornbury, Natalie Pedersen, Stephanie Fox, David Moore, Gemma Escott, Cathrine Bie Petersen, Hans Jakob Larsen, Rebecca Giles, Paul Stafford Allen, Ingo Bastisch

## Abstract

The ReAct (Recovery, Activity) project is an ENFSI (European Network of Forensic Science Institutes) supported initiative comprising a large consortium of laboratories. Here, the results from more than 23 laboratories are presented. The primary purpose was to design experiments simulating typical casework circumstances; collect data and to implement Bayesian networks to assess the value (i.e., likelihood ratio) of DNA results given activity level propositions. Two different experimental designs were used to simulate a robbery, where a screwdriver was used to force a door or window. Propositions and case information were chosen following laboratory feedback listing typical casework circumstances (included in the paper). Whereas the proposition representing prosecution’s view is applicable to many cases: the defendant forced the door/window with his screwdriver; the proposition representing defence’s view are more variable depending on what is alleged to have happened. In a direct transfer experiment, the defendant owned and used the screwdriver, but he did not force the door/window in question. An unknown person used the defendant’s stolen screwdriver. In an indirect transfer experiment, the defendant neither owned, saw, nor used the screwdriver, nor did they force the door or window. Instead, an unknown individual—someone with whom the defendant had interacted (e.g., by shaking hands or touching the same objects)—used the recovered tool to force entry. In the first experiment, if the defendant forced the door/window, then s/he is the last handler of the screw driver, whereas if the tool was stolen and used by an unknown person, the defendant is the first handler. For the second experiment, given the defence view, the defendant never held the screwdriver. We envisaged the situation where an object manipulated by the defendant (or the defendant himself/herself) would be touched by the unknown offender who would then force the window. The time delay between the touching of the object and the forcing of the window was 0, 1h, and 2h. This was accomplished by handshaking experiments, where the person forcing the door would shake a person’s hand, and then use the screwdriver either immediately after hand-shake or after 1, 2 hours. It was found for the direct transfer experiment that unless a single contributor profile aligning with the known person’s of interest profile was retrieved, the results did not allow to discriminate propositions. On the other hand, for the indirect transfer experiment, both single and major contributor profiles that aligned with the person of interest (POI) supported the proposition that the person used the tool rather than an unknown person who had touched an object, when indeed the former was true. There was considerable variation in median recoveries of DNA between laboratories (between 200pg - 5ng) for a given experiment if quantities are taken into account. These differences affect the likelihood ratios given activity level propositions. More than 2,700 samples were analysed in the course of this study. Two different Bayesian Networks are made available via an open source application written in Shiny R: `Shiny_React()`. For comparison, all datasets were analysed using a qualitative method categorised into absent, single, balanced or major given contributors.

The need for standardisation of methods is highlighted, along with the need for new methods to assess probability of laboratory dependent DNA recovery. Open access databases that are made freely available are important adjuncts

## 1. Introduction

There are two main categories of papers in the literature that explore DNA transfer: those that characterise some aspect of DNA persistence, transfer, and recovery, without interpreting the DNA findings given activity level propositions. And those attempting to assign likelihood ratios, for example by building models with Bayesian networks, usually with some specific case in mind, along with relevant data. Many different methods have been published, but there is no agreed standard to collect or interpret data (see earlier discussions by [1, 2]). Whereas there is much greater awareness of the limitations of DNA profiling evidence given sub-source propositions, the formal evaluation of DNA transfer issues, such as through the use of Bayesian networks, remains relatively rare in court proceedings. Although the theory is well developed; implementation lags behind. There are a number of reasons for this: a) the theory is complex b) laboratories lack training c) law enforcement and court are unaware that formal assessment can be done d) there are a lack of data to help inform probabilities. When a scientist is asked to help a court, it is inevitable that questions will be raised about ’how’, ’why’, ’when’ a DNA profile became evidential. If relevant data exist in the literature, then a scientist may cite this as evidence to provide the value of support given an activity level proposition, either in verbal or in quantitative terms. However, in doing so, *there is an implicit assumption that presupposes reproducibility of results between laboratories*. In the absence of such evidence, there is an inevitable leap of faith involved when a laboratory simply adopts findings from another, without corresponding experimental verification.

At this juncture, it is worthwhile drawing a parallel with methods used to interpret DNA profiles, given sub-source propositions. This field of forensic genetics is very well established. Laboratories follow a standard universal practice to sample local relevant populations and convert this data into frequency databases. These databases are used to calculate likelihood ratios. Even though genotype probabilities are very similar across different jurisdictions, for a given ancestral population, the standard practice is always for the laboratory to generate their own database, before reporting cases to court. The resources required to validate and accredit is not a trivial exercise.

Whereas investigations to prepare frequency databases is mainly *independent* of the methodology used (for a given set of loci and multiplex), this is not true for methods that consider activity level propositions, because interpretation depends upon the recovery of DNA profiles from contributors and this is function of many factors. A non-exhaustive list is:

1. The sampling method used
2. Extraction method
3. The quantification method
4. Quantity of extract forwarded to PCR
5. The PCR method
6. CE method / settings
7. Genotyping methods / thresholds
8. Shedder status of contributors

The question that inevitably follows is whether the same effort required to validate a method given sub-source propositions should be applied to methods given activity level propositions? Can laboratories rely upon data from external sources instead?

Over recent years, there has been much more awareness about reporting the value of DNA results considering the alleged activities, and court-attending scientists often have to give evidence to help address this issue. To do this, they may resort to the body of literature in order to make generalisations, and there are many examples of this - for reviews see [3, 4]. To date, there has been limited work to investigate reproducibility of DNA recovery between laboratories, although [5, 6, 7], examined this across 2-4 laboratories. Leuenberger et al [8] compared extraction methods across multiple laboratories in relation to drug testing.

The difficulty is compounded by lack of data available for independent scrutiny; the receiver of the information has to rely upon isolated peer reviewed material.

In general, published papers primarily speak to experiences of a particular laboratory. This does not mean that these same experiences are applicable to different laboratories, especially those that run different systems and processes.

### 1.1. Aims

The ReAct project, was instigated as a collaborative exercise to generate a body of data from 23 laboratories (plus some labs with partial results) with the following objectives

1. To identify two kinds of cases, typically encountered in casework
2. To standardise experimental protocols for laboratories to simulate specific case circumstances
3. To apply the method to obtain results from participant laboratories
4. To generate an open-access database of results
5. To produce open-source software that can form the basis of a repository for a collection of Bayesian Networks
6. To examine reproducibility between different laboratories
7. To gain experience with coordinating large projects with multiple participant laboratories and to describe the associated challenges

### 1.2. Experiments undertaken

Data were collected for cases that simulated a burglary using a screw-driver. Three separate experiments, described in section 2, were undertaken to determine quantities of DNA recovered from first and second handlers of the screwdriver, along with indirect transfer after handshake.

### 1.3. Results generation and analysis

The results were collated and analysed with an R application, ShinyRFU(), that utilises EuroForMix, to determine mixture proportions (*M*_*x*_) of contributors. Each laboratory determined quantity of DNA recovered; combined with *M*_*x*_ per contributor, DNA recovery per contributor is calculated. These values were used directly in Bayesian networks to compute likelihood ratios. A package written in Shiny R https://cran.r-project.org/web/packages/shiny/index.html called ShinyReact() has been made available, along with all data collected from 23 laboratories. These data were used to inform Bayesian networks for the latter two experimental designs described previously. Users of the software are able to choose a dataset to analyse, define the input Bayesian network parameters and calculate likelihood ratios for their findings, along with bootstrap derived confidence intervals. A user manual is available. Results are also compared to a qualitative method based on mixture proportions without DNA recovery attribution.

A huge amount of data has been collected. In fact, a total of 2,700 DNA profiles have been analysed and all these data are available for independent scrutiny (see resources section 2.22).

This work represents a massive undertaking without any parallel in the literature. The main purpose is to summarise the current work and to describe resources available. The project has identified that further work is needed before firm recommendations on the way forward can be made. Since data are made freely available, the community is free to carry out independent assessments.

Whereas raw genetic data cannot be made available because of privacy rules, all of the derived and associated data e.g., DNA quantification results and mixture proportion (*M*_*x*_) values used in this analysis are made available (see resources section 2.22) for independent review and study. Data collections and programs will be subject to continual addition and changes, and are subject to version control.

The paper is structured as follows:

1. The materials and methods section details the statistical methods used in the analysis, along with the location of resources.
2. Formulation of propositions based on a review of typical casework from laboratories.
3. Trends were analysed in a number of ways:
  a. Analysis of median quantities of DNA recovery for each class of contributor analysed, along with ranges.
  b. The use of beta-binomial distributions to provide probability densities of recovery, in terms of binary presence/absence across all laboratories.
4. A comparison of continuous vs. qualitative models, where the latter takes no account of DNA quantity.
5. A risk analysis to investigate misleading evidence in *H*_*p*_ = *true* and *H*_*d*_ = *true* (ground truth experiments).
6. A discussion on whether generalisations of results can be made across laboratories, along with use of proxy experiments that can be used in general casework.
7. The way forward for future work/ analyses to be undertaken to address the challenges identified by the work undertaken.

## 2. Materials and Methods with examples

### 2.1. Experiment 1: Direct transfer by a single participant

Experiment 1 was a control experiment. A cleaned screwdriver was placed (by the researcher wearing appropriate protective clothing) on a clean sheet of paper next to where it was used. At least one hour after washing hands, a participant fastened a screw into a wooden block, the screw was loosened and tighten several times so that the handling time with the screwdriver lasted a total of 5 minutes. Both hands could be used. He/she then placed the screwdriver on a separate clean sheet of paper next to where it was used. Directly after the procedure was carried out, the handle of the screwdriver was sampled according to the laboratory’s casework protocol (swabbing or tape lift, whole or partial handle) and the methods used noted in a table. A “Supplementary Information” spreadsheet was used to record the exact time since hand-wash and the activity performed before sampling. The protocol was then repeated for all participants in the experiment with cleaned screw-drivers and a new (clean) screw. The experiment was repeated by 20 or more different participants per laboratory

### 2.2. Experiment 2: Direct transfer by two participants

For this experiment participants were paired into 10 or more sets. At least one hour after washing hands both participants in a pair fastened a screw into a wooden block (as for experiment 1); the screw was loosened and tightened several times. Both hands could be used. The procedure was as follows: The screwdriver was placed on a clean sheet of paper next to where it was used. The first participant (the first handler) picked it up, used the screwdriver (for a total of 5 minutes) and returned it to the same place it was picked up from. The second participant (second handler) then picked it up and handled the screwdriver in the same manner with a total handling time of two minutes. He/she then placed it onto a separate clean sheet of paper next to where it was used. Directly after the procedure was done, the handle of the screwdriver was sampled according to the laboratory casework protocol (as for experiment 1) and the methods used were noted (see resources section 2.22). The protocol was then repeated for all pairs of participants in the experiment with cleaned screwdrivers and a new (clean) screw.

### 2.3. Experiment 3: Secondary transfer experiment

The two participants in a pair shook hands for 1 minute (firm handshake to simulate social contact between two individuals). Purpose was to maximise transfer between the two participants. One of the participants (referred to as the last handler) then grasped the handle of a cleaned screwdriver, and fastened a screw into the wooden block; the screw was loosened and tightened several times, so the handling time with the screwdriver lasted a total of 2 minutes. The full procedure, with a new handshake was performed a total of three times (non-consecutively) with a time interval of 0, 1 and 2 hours between shaking hands and handling the screwdriver. Both participants in a pair acted as the handler of the screwdriver for all time intervals.

Directly after the procedure had been carried out, the handle of the screw-driver was sampled according to the laboratory casework protocol and the methods used noted. The protocol was then repeated for all pairs of participants in the experiment with cleaned screwdrivers and a new (clean) screw.

### 2.4. Datasets

The datasets were anonymised and given a code (Lab 1-Lab 23). For labs providing results from more than 1 multiplex the multiplex abbreviation is given in the lab name (e.g. `lab18_F6C` and `lab18_GOF`). Combined datasets of laboratories were also prepared. These are defined in supplementary material 1, along with laboratories comprising the category:

1. Lab 201: Experiment 2 compilation
2. Lab 500: Highest tertile of LRs obtained from experiment 2
3. Lab 100: Mid tertile of of LRs obtained from experiment 2
4. Lab 600: Lowest tertile of LRs obtained from experiment 2
5. Lab 300: Experiment 3 compilation

Data compilations for the above are available (see resources section 2.22 for links)

### 2.5. Calculation of mixture proportions with ShinyRFU()

In order to derive quantities of DNA per contributor, DNA profiles were analysed using EuroForMix [9, 10], which outputs mixture proportions (*M*_*x*_) per contributor. The process was automated using the the ShinyRFU() program (see resources section 2.22). The ShinyRFU() program receives a script which lists the reference and ’evidence’ sample results, along with a hypothesis file that lists the number of contributors and the propositions to consider. An example, along with a user manual is provided (resources section 2.22) and described by Gill et al [11]. Mixture proportions were estimated. For any given sample result there are three categories to take into account:

1. The first handler (FirstH)
2. The second or last handler (LastH)
3. Unknown contributors (*U*)

The propositions considered here to estimate the mixture proportions are necessarily at sub-source level. N is the number of assigned contributors. The results were considered given the following propositions:

*H*_1_: The DNA is from first handler, the last handler (and (N-2) unknown)

*H*_2_: The DNA is from first handler and (N-1) unknown. We defined the results *E* as:

- *G*_*s*_: The DNA profile of the trace
- *G*_*f*_ : The DNA profile of first handler
- *G*_*l*_: The DNA profile of last handler

*I* is the information used regarding the model and the relevant population.

The likelihood ratio equation used in the ShinyRFU() calculation that considers sub-source propositions was as follows:

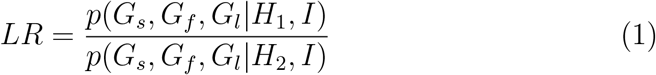

We used the ground truth experiments where the person used the tool to estimate *M*_*x*_ values for the first handler, the last handler and the background (i.e., unknown contributor(s)). The LR in equation 1 can be utilised as an indicator of quality of the profile and is listed in the compilation of data. Note that the first handler, FirstH, is conditioned under both *H*_1_ and *H*_2_, and does not necessarily reflect the value of the comparison that would be reported in casework when considering sub-source propositions. This is because either the first or last handler may be unknown.

### 2.6. Quantification of various contributors to a DNA profile

We can generalise that every DNA profile that is used in evaluation consists of a set of three contributors that is embodied in the experimental design:

- An individual who was the last person to handle the tool as the ’perpetrator’, abbreviated to ’LastH’ in the text (all experiments)
- An individual (abbreviated to ’FirstH’ in the text) who has either previously handled the tool (experiment 2), or shook hands with the last person to handle the tool
- Unknown individual(s), who under *H*_*p*_ is classed as background, but under *H*_*d*_ may also include the perpetrator (direct transfer)

Within our experimental design, since FirstH and LastH are known individuals, we can assign quantities that are recovered to all three categories in the set. In real case work, dependent upon the propositions, one handler may also be unknown, but we can nevertheless use our data to inform expectations. For expediency, and ease of calculation, we combine the quantities of all unknown contributors together.

### 2.7. Calculation of DNA quantities recovered

A number of different quantification methods were used by laboratories (recorded in ”information sheets” in excel files - see resources). No attempt was made to assess differences between quantification methods in this paper.

DNA quantities per *μl* (*Q*) were recorded by the laboratory. To convert into total recovery (*Q*_*T*_), the value was multiplied by the elution volume (*E*_*v*_)

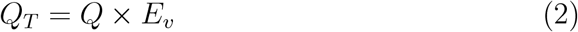

Then the proportion of DNA attributed per contributor was calculated by multiplying by *M*_*x*_; e.g., for the last handler (LastH):

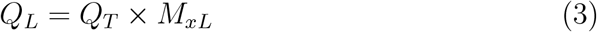

The calculation was repeated for first handler and unknown contributors. To simplify calculations, DNA quantities of unknown contributors are combined

After adjusting for dilution factors [11], RFU values, can be converted into DNA quantities. There is a log-linear relationship between the two variables (Figure 1). In this example, the data-set exhibits a robust fit with an R-squared value of 0.98. These analyses were repeated for all datasets; not all fits were as good (results for all analyses are available from the ShinyReact output)

**Figure 1:**
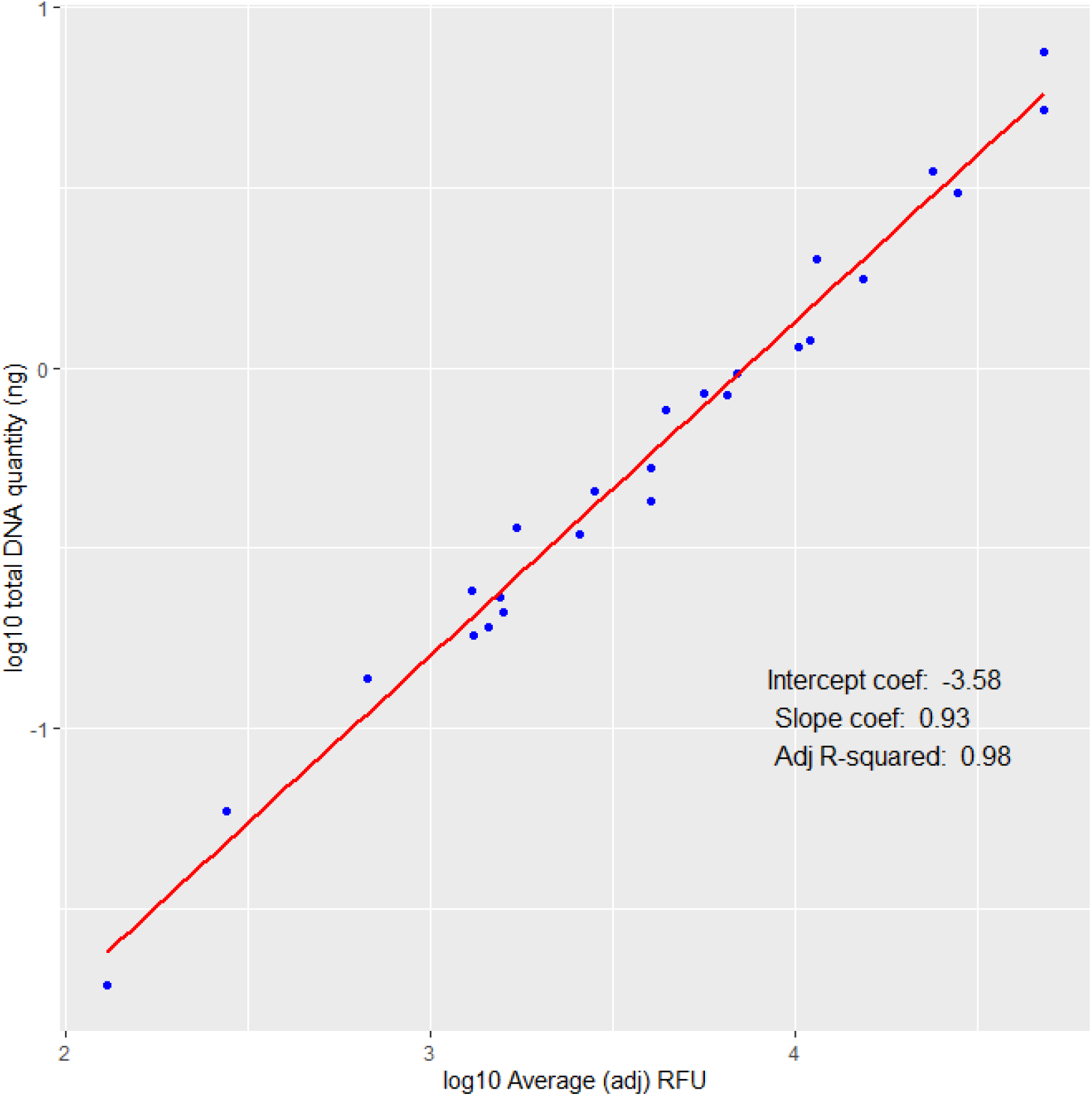
Log-log regression of DNA quantity vs. RFU from expt 1 data

### 2.8. Median polish to analyse trends of DNA recovery

Median polish [12] is a robust statistical technique used primarily for exploratory data analysis and to simplify the interpretation of multi-way tables, especially in the context of analyzing additive models. It is particularly useful for handling data with outliers or non-normal distributions. Here, median polish was used to decompose the matrix of DNA quantities into an additive model where the observed data is expressed as the sum of row effects, column effects, and an overall median. This decomposition helps in understanding the underlying structure of the data. Unlike mean-based methods, median polish is less sensitive to outliers. This robustness makes it suitable for datasets where extreme values might distort the analysis if traditional methods were used.

The R program ”medpolish” was used for all calculations.

Recovered quantities of FirstH, LastH and Background DNA were log10 transformed: a small value of 0.001ng was added to each datum prior to transformation, to obviate zero values. Full details of the analysis are available in Supplement 4

### 2.9. A compilation of case-work examples to help decide experimental design and propositions

Laboratories were asked to send examples of typical casework circumstances, along with their respective propositions that were used in evaluative reports. These examples are collated, discussed and summarised in Supplements 2 and 3. The decision of the experimental design and associated activity propositions that are followed in this paper are based upon these reports, but generalised.

### 2.10. Experiment 2 case circumstances and propositions

#### 2.10.1. Outline of the case circumstances

A tool (screwdriver) is used to force open a door in a burglary. The tool has been left at the crime scene, and there is no doubt that it was used by the perpetrator.

Suspect X (a known individual) is arrested and accused of the crime. He states that it is his tool, but that it had been recently stolen, and that he did not force the door in the burglary.

#### 2.10.2. Findings

The DNA aligns with X. The value of the comparison given sub-source level propositions is assigned. It is assumed that it is not disputed that DNA is from the suspect hence we do not consider this further. There may or may not be DNA from an unknown contributor(s) present.

#### 2.10.3. Propositions

1. *H*_*p*_: Mr X (known individual) forced the door with his own tool.
2. Or *H*_*d*_: An unknown person forced the door with Mr X’s tool.

It is not disputed that the tool belongs to Mr X. The analysis evaluates the probability of the evidence if the suspect was either the last handler (*H*_*p*_) or the first handler (*H*_*d*_) of the tool.

### 2.11. Experiment 3 case circumstances and propositions

#### 2.11.1. Outline of the case circumstances

The outline of the case circumstances are the same as described in the previous section (experiment 2). The difference is that the suspect denies all knowledge of the screwdriver and states that he/she has never seen or held it before. He had nothing to do with the incident.

#### 2.11.2. Findings

Same as in the previous section reiterated here: The unknown DNA profile aligns with Mr X’s DNA profile. It is assumed that it is not disputed that DNA is from Mr X, hence we do not consider this further. There may or may not be DNA from an unknown contributor(s) present.

#### 2.11.3. Propositions

Same as in the previous section:

1. *H*_*p*_: Mr X (known individual) forced the door with his tool.
2. *H*_*d*_: An unknown person - who frequents the same premises as Mr X - forced the door

The analysis evaluates the probability of the evidence if the suspect was either the last handler of the tool (*H*_*p*_) or there was some kind of unknown social activity with an unknown individual which we simulate by handshaking (*H*_*d*_).

### 2.12. Statistical formulation

There are two Bayesian networks that address experiments 2 and 3 respectively. They are based upon the Bayesian network described by the ISFG DNA commission (see supplement 1 of [13]). In order to enable rapid computation, and to ensure that the method was open source (i.e. free of any commercial software restrictions), formulae were raw programmed into `Shiny_React()`.

### 2.13. Nomenclature

There are four probabilities used to inform the Bayesian network denoted by *s, t, t*^*′*^, *b* to calculate probability of respective events: FirstH, LastH and Background

1. *s*: FirstH, either probability of recovery of first handler (experiment 2) or probability of secondary transfer (experiment 3).
2. *t*: LastH, the probability of direct transfer from the last handler. This probability is exclusively assigned under *H*_*p*_.
3. *t*^*′*^: LastH, the probability of direct transfer from an unknown contributor under *Hd*.
4. *b*: Background: Probability of background from an unknown contributor(s). This probability is assigned under both *H*_*p*_ and *H*_*d*_.

### 2.14. Mixture proportion (Mx) categorical method

In each DNA profiling result, there are three possible contributors: last handler, first handler, unknown contributor. Each of these can be categorised into a mutually exclusive outcomes: Absent, Single, Balanced, Major. This is similar to the categories used by [14], except we refer to ”Balanced” rather than ”mixed DNA profile with no major”.

The precise definitions of the various categories are rule-based and coded into an R program:

1. If the contributor is present such that *M*_*x*_ ≥ 0.99 then the outcome is ”Single”
2. If the contributor is present such that *M*_*x*_ ≥ 0.7 and *M*_*x*_ *<* 0.99 then the outcome is ”Major”
3. If the contributor is present such that *M*_*x*_ *>* 0.01 and *M*_*x*_ *<* 0.7 then the outcome is ”Balanced”
4. If the number of contributors (NOC=0) then the outcome is ”Absent”
5. If the contributor is not observed: *M*_*x*_ ≤ .01, then the outcome is ”Absent”

This is the ’basic’ qualitative (*M*_*x*_) model, because it does not take account of other factors such as the DNA amount (total and per contributor), or numbers of alleles recovered. Consequences are discussed further in the text.

The POI is either: the last handler (LastH) or the first handler for experiment 2; or the contributor whose profile was secondarily transferred (FirstH) in experiment 3; or an unknown contributor. The probabilities of outcomes are assigned to each class of contributor as shown from the example in Table 1 (with conditioning shown in Table 2).

**Table 1:** Example of a probability table, experiment 2, lab 4 NGM. LastH and FirstH are last and first handlers respectively.

| POI category |  |  |  |  |
| --- | --- | --- | --- | --- |
| Experiment | Category | LastH | FirstH | Unknown |
| Ex_2 | Absent | 0.06 | 0.19 | 0.44 |
| Ex_2 | Single | 0.19 | 0.06 | 0.03 |
| Ex_2 | Balanced | 0.39 | 0.72 | 0.50 |
| Ex_2 | Major | 0.36 | 0.03 | 0.03 |

**Table 2:** Probability assignments from Table 1 example.

| Condition | s | t | t' | b |
| --- | --- | --- | --- | --- |
| POI is single contributor: | Single \| FirstH | Single \| LastH | 1-Absent \| LastH | NA (cancels) |
| POI is balanced contributor | Balanced \| FirstH | Balanced \| LastH | Balanced \| LastH | Balanced \| Unknown |
| POI is major contributor | Major \| FirstH | Major \| LastH | Major \| LastH | Major \| Unknown |

Note that for a single contributor, there is no unknown contributor, hence the assignment 1 − *t*^*′*^ is required under *H*_*d*_ (the probability of not observing an unknown) where *t*^*′*^ = 1 − *Absent*

### 2.15. Continuous method

The method differs from the *M*_*x*_ categorical method in that log normal probability distributions of recovered DNA quantity per contributor are fitted as described by [15]. Contributors are in the classes (LastH, FirstH and Background),as described for the *M*_*x*_ categorical method.

#### 2.15.1. Qualitative step

There are two steps in the continuous model. The first is qualitative. For each contributor per sample, presence of recovered DNA was recorded based upon the total quantity per contributor *Q*_*L*_ *>* 0.001*ng* (eq: 3) and *k* is the proportion of samples where the contributor is absent.

#### 2.15.2. Calculation of probability distributions

In the second step, a dataset comprising *Q*_*L*_ *>* 0.001*ng* was compiled and analysed following supplement 2 in [15]; summarised here:

1. Log-normal distributions were fitted to each dataset, using the fitdis-trplus() package in R. Diagnostic checks were made to ensure the data were a reasonable fit. Example plots are provided in (Fig. 2).
2. There are two log-normal parameters *μ* and *σ*, where *x >* 0: *μ* is the location parameter and *σ* the scale parameter of the log-normal distribution. The third parameter *k*=the proportion of observations where no DNA was recovered (from the first step).
3. Cumulative probability (1-cdf) functions were fitted, scaled to *k*. An example is shown in fig: 2 for each each contributor per experiment. For a given quantification result, the probability of observing a quantity of DNA greater than the observed value is assigned.

**Figure 2:**
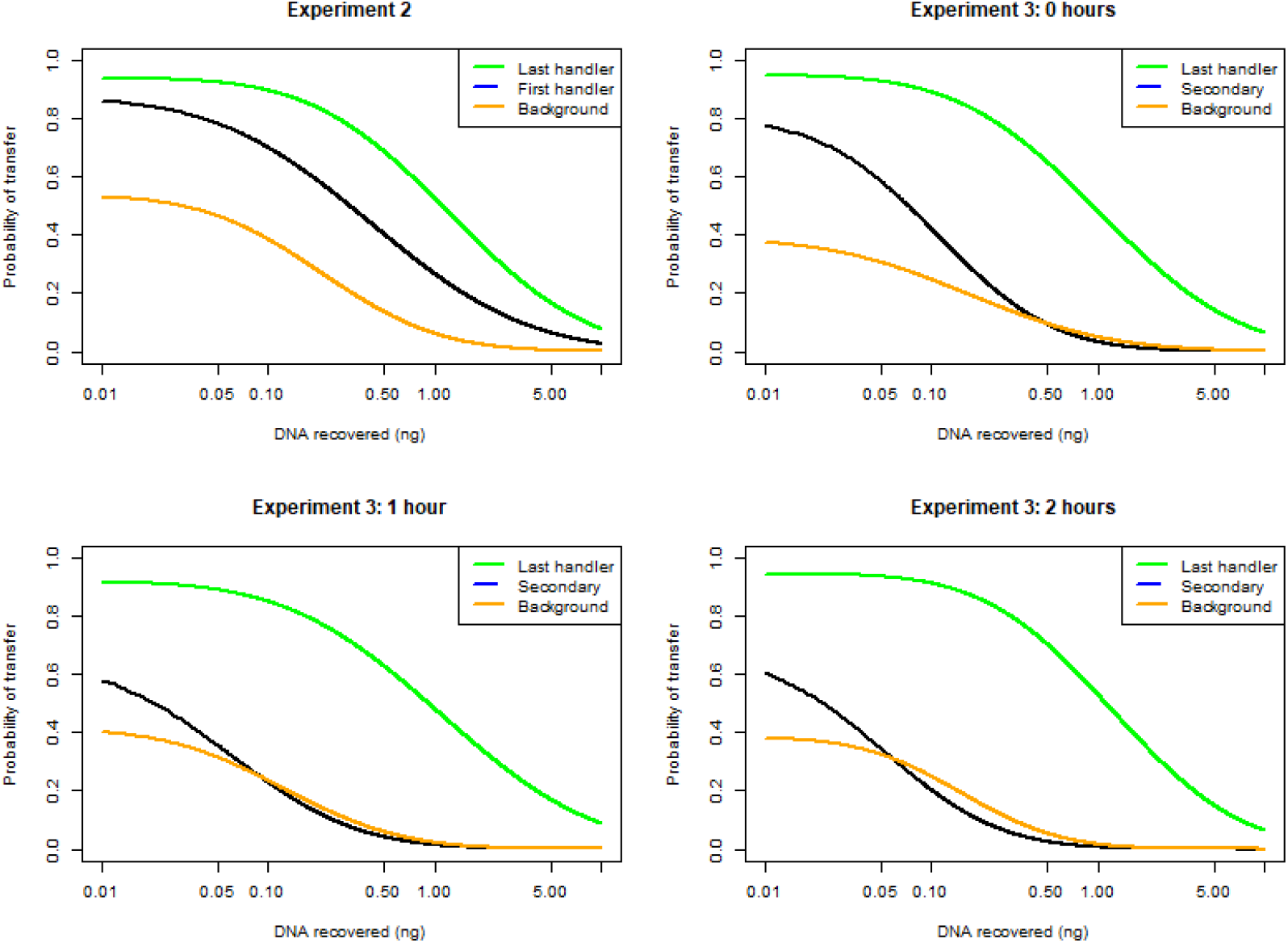
DNA Quantity recovered vs Probability of transfer plots for experiment 2 and 3, from lab 201 (compilation of laboratories) results.

### 2.16. Probability density assignment of presence/absence of contributors: categorical method

Beta-binomial Bayesian models are used when data are categorised. To help understand the foregoing, the reader is recommended to familiarise him/herself with the concepts by reading one of the many guides available from educational sources. The following is recommended as a gentle introduction to the subject: https://www.bayesrulesbook.com/chapter-3

Following the rule-set in section 2.14, data can be subdivided, per laboratory, according to categories that count the presence/absence of LastH,

FirstH and Background/Unknown contributors (listed in Supplements 5A and 5B). Count data are converted into observed proportions per contributor per laboratory. The means and variances are calculated across data sets. Probability densities are found by applying beta-binomial modelling. The probability density is described by two parameters: alpha and beta. Mean probabilities and variances can also be derived directly from these parameters. The method of moments was used: program written in R-code. These distributions give a compact overview of the prior expectations of probability densities of presence/absence of contributors across all laboratories and are a useful adjunct to the analysis of medians that were described previously.

### 2.17. LR formulae

The next step is to calculate likelihood ratios from the Bayesian network, using the formulae outlined below:

### 2.18. Experiment 2

#### 2.18.1. ’POI only’ observed

Background cancels in the formula

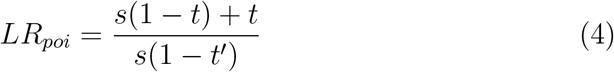

#### 2.18.2. POI and unknown contributor(s) observed

Background does not cancel

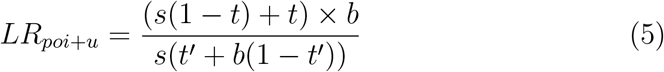

### 2.19. Experiment 3

The formula is the same as for experiment 2 (eq4, except that *s* = 0; therefore the first part of the expression does not appear. The denominator is unchanged.

#### 2.19.1. POI only

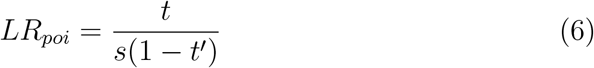

#### 2.19.2. ‘POI and unknown’ is observed

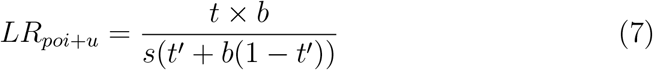

### 2.20. Avoiding probabilities of one and zero

Following on from the introduction in section 2.16 to beta binomial Bayesian modelling, to carry out calculations, a beta prior *a* = *b* = 1 is adopted, so that distribution *π*(*θ*) = 1, 0 *< θ <* 1, is uniform (called the ”principle of insufficient reason” by Laplace). Also known as an ”uninformative” prior: the prior mean = 0.5 for the presence/absence method used in the continuous model. The posterior mean estimator, known as Laplace’s estimator, is:

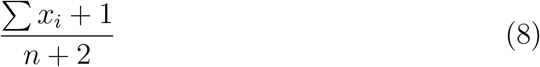

where *x* is an observation and *n* is the sample size. For the *M*_*x*_ categorical method, results are divided into four different categories; Absent, Single, Balanced, Major. Since there are now four categories, instead of two (i.e. for each category the prior = 0.25), this is taken into account as follows:

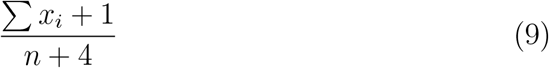

Posterior estimators have the useful property of avoiding situations where either there are no observations, or conversely, an observation in every sample. Without the Bayesian estimator, this results in *Pr* = 0 or 1, which would result in mis-calculations. If *s* = 0 then the LR cannot be calculated in any of the equations as it results in an infinite value. Similarly, in eqs. 4,6, if *t* = 1 then 1 − *t* = 0, hence LR= infinity. Both are common occurrences in the datasets analysed.

### 2.21. Qualitative LR based on analytical threshold

Laboratory specific analytical thresholds (*AT*_*RF U*_) are based upon RFU measurements. However, we use quantification instead, in order to standardise a method that does not depend on peak heights. An equivalent DNA quantity per laboratory *AT*_*Q*_ is calculated by regression analysis as described in section 2.7. Since there are two alleles per locus, the equivalent *AT*_*Q*_ is approximated from 2 *× AT*_*RF U*_. Users can calculate qualitative LRs, based on this presence/absence threshold in the Shiny React program.

### 2.22. Resources available

#### 2.22.1. Supplementary material

1. Supplement 1: A list of laboratories used in combined data-sets (section 2.4)
2. Supplement 2: Compilation of case examples submitted by participating laboratories (section 2.9).
3. Supplement 3: A summary spreadsheet of case examples submitted by participating laboratories (section 2.9).
4. Supplement 4: Analysis of median recoveries from laboratories (section 3.1).
5. Supplement 5 part A: Beta Binomial distribution analysis (section 3.4).
6. Supplement 5 part B: Beta Binomial distribution analysis (using sub-source LR*>*1000 as a filter criterion). (section 3.4)
7. Supplement 6: Analysis of data using the *M*_*x*_ categorical method (section 3.6.4).
8. Supplement 7: The ’at risk :*H*_*d*_ = *true*’ database (section 3.11).
9. Supplement 8: *H*_*p*_=true analysis (section 3.13).

#### 2.22.2. Online resources

The main online resource is https://sites.google.com/view/altrap/ enfsi-react-project. This website has links to all of the online resources which includes:

1. A summary of the ReAct project.
2. Links to the `Shiny_React()` program and user manual.
3. ShinyRFU() program access and user manual available at https://sites.google.com/view/altrap/average-rfu-method.
4. Links to excel spreadsheet databases:
  a. `Grand_compilation` of all data compiled into a single excel spread-sheet.
  b. Laboratory specific folders containing data and the complete analysis using the `Shiny_React()` program.

Note that all analyses in this paper were carried out using version 2 of programs and data-sets.

## 3. Results and discussion

A challenge of the experiment was to compare the data from many different laboratories that use diverse systems; to recap:

1. Different sampling techniques.
2. Different extraction methods.
3. Different elution and PCR volumes.
4. Different multiplexes and PCR strategies.
5. Different analytical thresholds

All of the above affect the output that is visualised. The only thing in common to all laboratories is quantification (which is universally applied). However, even here, there are differences that include: a) different manufacturers b) different standards, i.e., it is possible that comparability of quantification and associated standards, are themselves not standardised, and therefore subject to error and unpredictable variation between laboratories.

### 3.1. Analysis of medians

Median recovery rates in *pg* were calculated per contributor: LastH, FirstH and Background, for each experiment. The full table is available in Supplement 4. An abbreviated version is shown in Table 3. Laboratories are ranked by the ’LastH experiment 1’ column. It can be seen that there is considerable variability in recovery rates per laboratory, ranging from 200pg to 5ng. Low LastH recoveries usually result in low recovery rates for FirstH (e.g. labs 10,12,6). The median recovery rates across all laboratories are also shown, e.g., LastH, experiment1 = 1080pg.

**Table 3:**
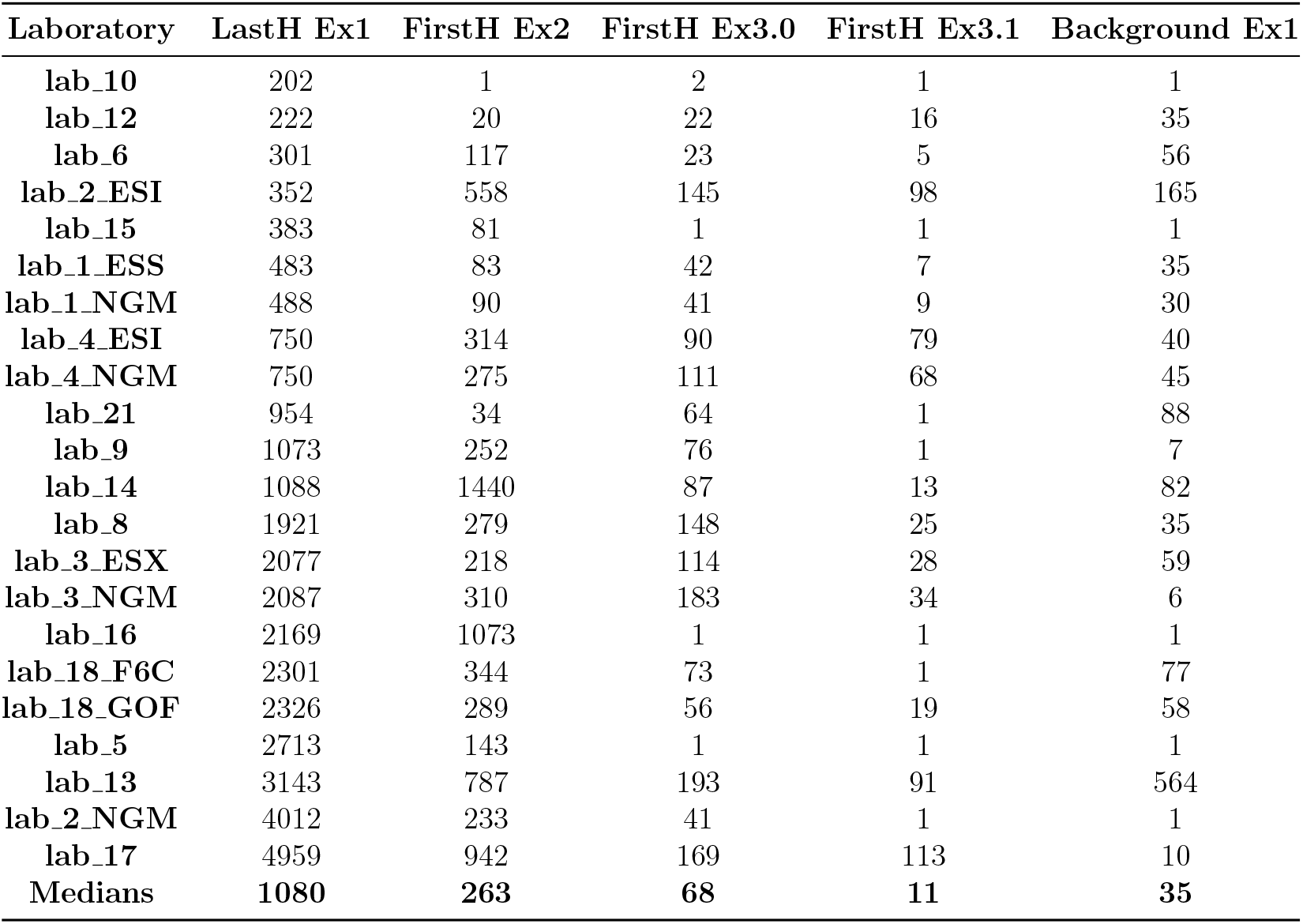
Tabulated raw medians of selected experiments and LastH, FirstH and background recoveries in pg. Ranked by LastH, experiment 1. The median value of medians is shown in the final row.

Further analysis to place findings into context, follows in the next section.

### 3.2. Median polish to examine trends of DNA recovery per contribution type

A two-way table represents the relationship between the response variable (the quantity of DNA recovered), *y* and the two ’effects’: the experiment effect *α*_*i*_ (a total of 14 categories shown in Table 4) and the laboratory effect *β*_*j*_. The residuals *ϵ* represent the portion of quantity recovery that cannot be explained by either factor. *μ* is the common value. This is the overall median DNA recovery across all labs and all experiments. The model is described as:

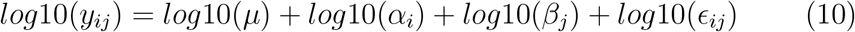

**Table 4:** Median polish experiment (row) effects *log*10(*α*_*i*_) in *ng* are shown and the detransformed results 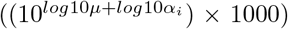 where *log*10*μ* = −1.29. For the detransformed data (*pg*), black type refers to LastH, red type to FirstH and green type to Background recovery.

| Experiment | Effects | Detransformed (pg) |
| --- | --- | --- |
| LastH_Ex1 | 1.52 | 1685 |
| Background_Ex1 | -0.12 | 38 |
| LastH_Ex2 | 1.32 | 1053 |
| FirstH_Ex2 | 0.74 | 278 |
| Background_Ex2 | -0.51 | 16 |
| LastH_Ex3.0 | 1.25 | 896 |
| FirstH_Ex3.0 | 0.12 | 67 |
| Background_Ex3.0 | -1.41 | 2 |
| LastH_Ex3.1 | 1.33 | 1086 |
| FirstH_Ex3.1 | -0.55 | 14 |
| Background_Ex3.1 | -1.10 | 4 |
| LastH_Ex3.2 | 1.40 | 1278 |
| FirstH_Ex3.2 | -0.55 | 14 |
| Background_Ex3.2 | -1.12 | 4 |

The detransformed results in Table 4 strip out the laboratory effect and summarise median (*pg*) DNA recovery rates across all laboratories. In accordance with expectations, greatest recoveries are obtained from LastH donors: total recovery between 896-1685pg with a mean value=1.2ng.

For experiment 2, FirstH: the median value of 278pg was obtained. The second handler (LastH) physically removes first handler DNA. Consequently, the availability of FirstH DNA is reduced by c. 75%.

The recovery of FirstH DNA for experiment 3 was much lower. When samples are taken immediately after the handshake, a median of 67pg was recovered. In contrast, total DNA recoveries for one and two hour experiments were the same (14pg), which would be challenging to detect for the majority of laboratories.

It can be concluded that in our experiments, secondary transfer occurs with much lower probability than direct transfer. This has an affect on corresponding likelihood ratios, which is discussed in sections 3.6 and 3.7.

A comparison of expected median proportions of LastH, FirstH and Back-ground that are recovered per experiment across all laboratories are listed in (Table 5).

**Table 5:** Expected median recovery proportions calculated conditioned on the stated category, where the total DNA contributions are from LastH, FirstH and Background, except for LastH Ex1 where only LastH+Background are present.

| Category | Proportion |
| --- | --- |
| LastH Ex1 | 0.97 |
| FirstH Ex2 | 0.20 |
| FirstH Ex3.0 | 0.06 |
| FirstH Ex3.1 and Ex3. | 0.01 |
| Background Ex1 | 0.03 |

The low proportion of FirstH recovery in experiment 3.0 means that labs that recover low amounts of DNA will paradoxically record lower levels of secondary transfer. For example a total recovery of 300pg (lab 6) in 100ul elution volume where a total of 15*μl* is loaded for PCR contains an expected median 3pg, which is less than one cell equivalent and unlikely to be detected. A total recovery of 2ng (e.g. lab 3) results in 18pg loading of secondarily transferred DNA, which has a much better chance of detection, albeit as a low level mixture with LastH as the major contributor. Where secondary transfer has occcured after 1h+ since handshake, the proportion of recovered DNA drops to 0.01, further reducing the chance of detection.

### 3.3. Is a contributor present or absent? Success rates

Within the experimental design only, the identity of first and last handlers are known individuals as ground truth and consequently their presence is assigned Pr=1. We condition upon this in the calculation given sub-source propositions (equation 1) to calculate the mixture proportion, and this is used to assign DNA recovery per contributor. In actual casework, the ground truth is not known and presence/absence of a contributor is based upon laboratory criteria and it is not true that Pr=1, as in the experimental design. Before performing calculations of likelihood ratios (LRs) given activity level propositions, it is standard practice to first confirm that the LR given sub-source propositions is sufficiently high. This ensures that the comparison results support the proposition that a specific contributor, rather than an unknown individual, is the source of the DNA. [13]. However, there is no specification or guidance of what this threshold may be and it will vary between laboratories and may depend on case circumstances.

#### 3.3.1. A comparison of qualitative vs continuous models

Now, in casework, one contributor will usually be unknown and we do not know the ground truth of the known. Under these circumstances, as explained in the previous section, some extra information may be needed for the *qualitative* (*M*_*x*_) model before the presence/ absence of a contributor can be assigned in a binary way. For example, [14] used expert opinion to decide if a profile was interpretable based on *”the lack of the genetic information present on the EPG or because of the complexity”*. A limitation of such studies is that the criteria are undefined, and this has consequences for the adoption of a method by other laboratories (since they will use different criteria to define presence/ absence). This is alleviated by using strict rule-sets to decide presence/absence and future publications should always include such an assessment, otherwise how can other laboratories adopt findings? For example, a criterion could include a likelihood ratio given sub-source level propositions that is in excess of some arbitrary value. One could also account for the possibility that either it is DNA from the person or not, in our LR formula or Bayesian network.

Next we describe the fundamental differences between qualitative and continuous modelling to illustrate limitations of the former.

We have now established that a decision on whether the person’s DNA is present or absent is necessary for both models. If absent, then there is no second step; one could assess the value of this absence. However, if present, we move to the choice of how to describe the results and which model to choose. Both models are dependent upon data with ground truth observations. The qualitative (*M*_*x*_) model is categorical and dependent upon threshold criteria (such as *M*_*x*_ value, number of ’matching’ alleles etc.): the question that is asked by the qualitative model is *”What is the probability of DNA presence/absence of a contributor?”* The consequence is that a different model must be built for every set of criteria that laboratories utilise because the DNA presence/absence decision criteria are used to define it.

Continuous modelling does not suffer from this constraint. This is because the model is constructed from DNA quantities recovered from ground truth datasets: the question is very different: *”What is the probability of a given DNA recovery (in nanograms) if the POI handled the screwdriver?”*, regardless of whether DNA aligning with the POI is observed or not. If no DNA is recovered, this is integrated into the model and is nicely illustrated in Fig: 2 where the reader will note that the probability of (1-absence) of DNA (absence defined here as *<* 1*pg*) is shown on the y-axis: DNA from first contributors and unknowns are less likely to be observed in all cases.

Since the continuous model is entirely dependent upon ground truth observations to construct prior log normal distributions based on quantity of DNA recovered, the probability of observing DNA aligning with a contributor at a given DNA quantity is calculated. Unlike the qualitative model, the decision criteria of whether to report a case based on profile quality has no effect on the LR given activity level propositions. If the quality is too poor then the calculation is simply not made in the first place.

#### 3.3.2. Summary of important findings: adoption of models

In the foregoing we assume that laboratory A and B follow the same protocols, but the criteria to decide presence/ absence differ. What are the consequences if laboratory A wants to adopt the findings of laboratory B?

A single analysis is needed to create a log-normal distribution plot for the continuous model. The decision on whether to report the results based on activity level propositions is an entirely separate matter. It is valid for a laboratory to use broad presence/absence criteria when determining whether to report results, and this does not impact the model. Consequently, the model itself can be shared with laboratories that follow the same procedures but may have different interpretation guidelines.

For the qualitative model, a binary decision is made to decide presence or absence. Because these decisions currently tend not to be well defined, it inevitably leads to programming difficulties. Even if criteria are well defined, it still means that the model will need to be modified per adopting laboratory i.e instead of a single model (as described above), an extra layer of complexity is added, since the data will need reanalysis by laboratory B, according to their specific decision criteria. Of course, this can be solved if laboratories move away from vague to programmable criteria, and the next section shows how this can be achieved.

### 3.4. Probability densities using beta-binomial distributions

This section is only relevant to the qualitative (*M*_*x*_) method and continues the discussion about the crucial decision whether a contributor is present or absent. Rationale is described in methods section 2.16. Whereas the previous section 3.2 characterises raw recoveries (quantities) across laboratories, in this section, the distribution of recovery success rates across laboratories are examined. In the first instance, presence/absence of a contributor is decided with the ground truth approach described in section 3.3. Data pertaining to presence or absence of a contributor, measured per laboratory, are provided in Supplement 5 part A. The beta-binomial distribution was employed to model the likelihood of presence or absence across laboratories. The expected mean values per experiment 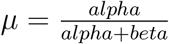 and their variances 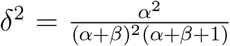 are listed in Table 6 and plotted in fig 3. The alpha and beta parameters are described as informative Bayesian priors which can be used instead of the flat priors described in section 2.20. In the previous section 3.3, we showed that a consideration of whether a contributor is actually present or absent is largely dependent on subjective criteria that are not standardised, and consequently practice is extremely variable across laboratories. Considerations may include the number of alleles aligning with the POI; a likelihood ratio given sub-source propositions of a specified magnitude [16]. It is also affected by the detection threshold applied and the number of contributors assigned by the laboratory. In this study, presence/ absence is defined by the rule-set described in section 2.14. This rule set was predicated by *M*_*x*_ *>* 0.01 as a lower threshold to define presence of DNA from a given contributor. We can call this the ’basic modelling assumptions’. Examination of the ”grand compilation” spreadsheet (see resources section 2.22) shows much variation in LRs given sub-source propositions, and if factored into the calculation, this will of course affect the perceived presence/absence of contributors^1^ along with a likelihood ratio given activity level propositions. To illustrate a possible casework alternative to the ground-truth conditioning used in the basic model, an additional criterion was added to the rule-set specifying that the LR given sub-source propositions must exceed a value of 1000: reanalysis shown in Supplement 5 part B. Results were similar. For example, the probability of observing the last handler was reduced to a mean of 0.93 (from 0.97) for experiment 1 and 0.85 instead of 0.96 for experiment 2 (variances were much larger, however). In the context of using the data to derive informative prior distributions, sensitivity analysis would be useful to determine the impact (which is likely to be small), but this exercise is reserved for future work.

**Table 6:** Parameters for the beta binomial, along with expected means and variance, calculated from 23 laboratories for all experiments.

| **LastH** | **Ex_1** | **Ex_2** | **Ex_3.0** | **Ex_3.1** | **Ex_3.2** |
| --- | --- | --- | --- | --- | --- |
| **alpha** | 9.37 | 4.58 | 2.45 | 1.3 | 1.51 |
| **beta** | 0.29 | 0.19 | 0.14 | 0.1 | 0.11 |
| **mean** | 0.97 | 0.96 | 0.95 | 0.93 | 0.93 |
| **variance** | 0.003 | 0.01 | 0.01 | 0.02 | 0.02 |
| **FirstH** |  |  |  |  |  |
| **alpha** | NA | 5.40 | 1.96 | 1.77 | 1.76 |
| **beta** | NA | 0.71 | 0.87 | 1.94 | 1.17 |
| **mean** | NA | 0.88 | 0.69 | 0.48 | 0.46 |
| **variance** | NA | 0.01 | 0.06 | 0.05 | 0.06 |
| **Background** |  |  |  |  |  |
| **alpha** | 1.49 | 1.22 | 0.98 | 0.83 | 0.74 |
| **beta** | 0.86 | 1.26 | 1.86 | 1.16 | 1.17 |
| **mean** | 0.63 | 0.49 | 0.35 | 0.42 | 0.39 |
| **variance** | 0.07 | 0.07 | 0.06 | 0.08 | 0.08 |

**Figure 3:**
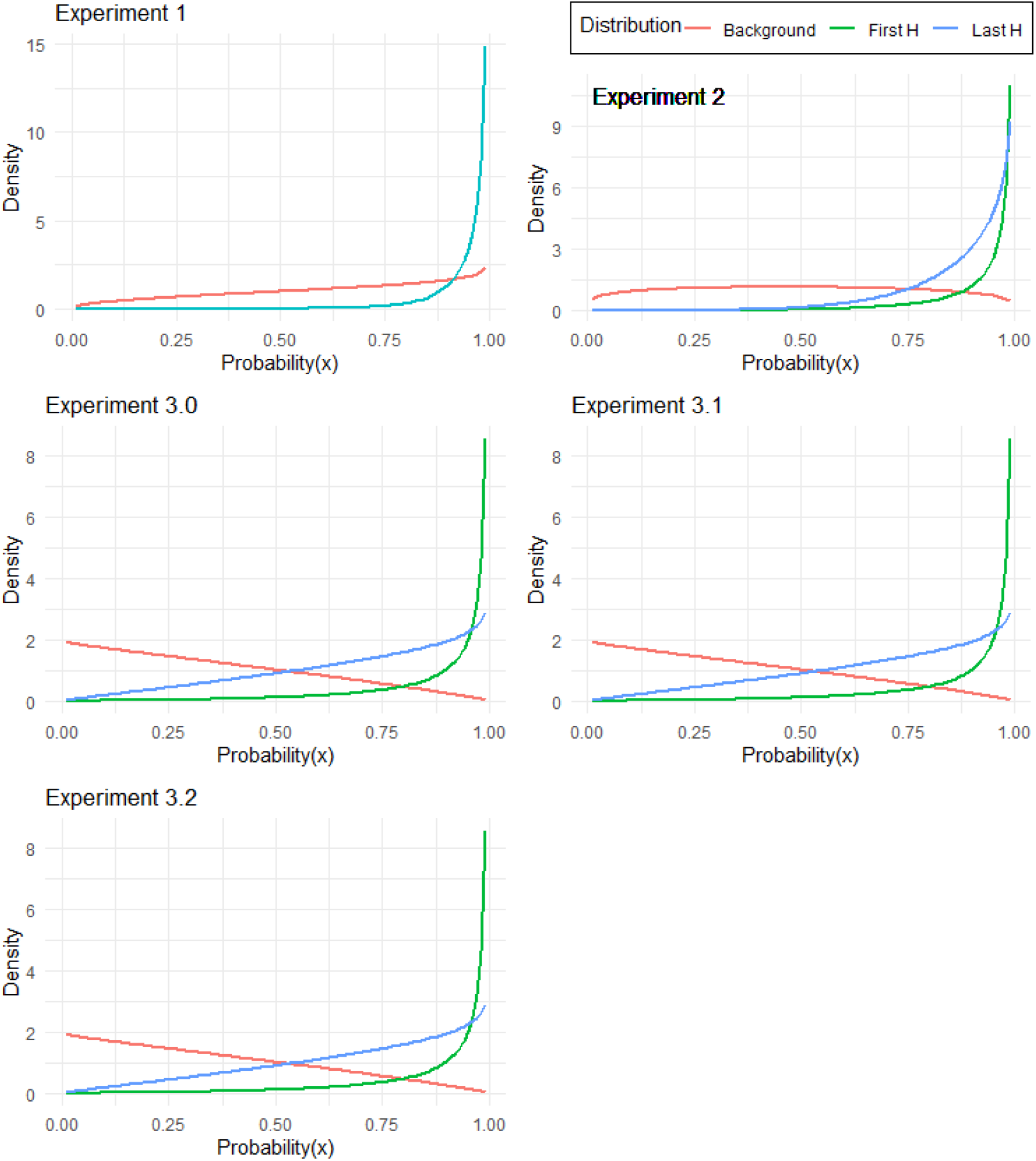
Beta binomial probability densities for presence of first and last contributors and background across all laboratories

Alternatively, if a particular laboratory wishes to derive Bayesian priors for their specific qualitative (*M*_*x*_) method, then this can be done with the existing dataset. As previously alluded to, there is no universal accepted method to decide presence/absence of a contributor. Within a Bayesian context, the purpose is to derive priors. Priors constitute our beliefs, and are not intended to be ’exact’, whilst recognising at the same time that priors can always be ’improved’ if more information is available.

In the foregoing discussion, we only consider the ground truth basic model in Supplement 5 part A and Table 6 where flat uninformative Bayesian prior distributions were used. The data show that when an individual is the last person to grasp a screwdriver as the last handler (LastH), similar results are obtained for all experiments. There is a mean probability *Pr >* 0.93 of DNA recovery (Table 6).

The mean recovery of the first handler for experiment 2 or handshaker (FirstH) for experiment 3 has a lower DNA recovery rate. For experiment 2, *Pr* = 0.88. For experiment 3, the success rate unsurprisingly depends upon the time since handshake: Experiment 3.0 (0 hours), *Pr* = 0.69; reducing to *Pr* = 0.46 for experiment 3.2 (two hours), with little difference for experiment 3.1 (one hour) at *Pr* = 0.48.

The probability of background recovery (i.e., from an unknown contributor) was much lower: highest for experiment 1 at *Pr* = 0.63; experiment 2 at *Pr* = 0.49; experiment 3 results ranged between *Pr* = 0.35 − 0.42. The distribution curves of background and FirstH are relatively flat and convergent in experiments 3.1 and 3.2.

In summary: LastH means and distributions are similar across all experiments. FirstH means are only slightly lower than LastH in experiment 2, and the consequence of this that LRs are low for mixtures (section 3.6). For experiment 3, the likelihood of DNA recovery from FirstH is dependent upon the time since handshake, hence the difference between FirstH and LastH is greater. This results in higher LRs (section 3.7).

Results of tests are compiled for all 23 participant laboratories. The `Shiny_React()` application is used to visualise the data and to carry out calculations (only for the continuous model). A description of the software is provided in the user manual (resources section 2.22).

### 3.5. Log normal distribution plots

Log normal distributions are plotted for each experiment per laboratory. An example is shown in fig. 2. Probabilities are used to calculate likelihood ratios in accordance with formulae provided in section 2.17. For each laboratory and for each experiment, results are both tabulated and plotted by the `Shiny_React()` application.

### 3.6. Experiment 2 LR results

#### 3.6.1. Continuous method

The median response across laboratories analysed from combined data of experiment 2 (lab 201) is shown in Table 7). If *H*_*p*_ = true, the *LR* increases as DNA recovery from the POI increases. There is a big difference in LRs for single vs mixed samples - support for the *H*_*p*_ proposition (*LR >* 20) was only achieved with single DNA profiles (i.e., non-mixed results).

**Table 7:** Experiment 2, lab 201 LR results. Continuous model results compared to Mx method. For the latter, the median LR is shown ”Across all labs” along with the results from all laboratories combined (lab 201). The results between the continuous and Mx methods can be directly compared. LRs shown in black type are ”Balanced”; red type ”Major” and blue type is defined as ”Single” with the Mx method, but is analysed as a mixture for the continuous method

|  | POI quant (ng) | 0.01 | 0.5 | 1 | 2 | 4 | 6 | 10 |
| --- | --- | --- | --- | --- | --- | --- | --- | --- |
| Single | 0 | 27 | 50 | 61 | 73 | 84 | 89 | 94 |
|  | 0.05 | 0.5 | 1.0 | 1.2 | 1.5 | 2 | 2 | 2 |
| Unknown | 0.1 | 0.5 | 0.8 | 1.0 | 1.2 | 1.4 | 1.5 | 2 |
| quants (ng) | 0.2 | 0.3 | 0.6 | 0.8 | 0.9 | 1.0 | 1.1 | 1.2 |
|  | 0.3 | 0.3 | 0.5 | 0.6 | 0.7 | 0.8 | 0.9 | 0.9 |
|  | 0.5 | 0.2 | 0.4 | 0.4 | 0.5 | 0.6 | 0.6 | 0.7 |
|  | 1 | 0.1 | 0.2 | 0.3 | 0.3 | 0.4 | 0.4 | 0.4 |
|  | 2 | 0.1 | 0.1 | 0.1 | 0.2 | 0.2 | 0.2 | 0.2 |
| Mx Category |  | Across all labs |  | Combined |  |  |  |  |
| Single |  | 55 |  | 191 |  |  |  |  |
| Balanced |  | 0.8 |  | 0.8 |  |  |  |  |
| Major |  | 2.6 |  | 4 |  |  |  |  |

For mixtures, the higher the proportion of unknown contributor, the lower the LR. Consequently, results either support *H*_*d*_ proposition, or are neutral. These same trends were observed in all participating laboratories (data for individual laboratories are available from the ”`Data_Output_v2`” folder - resource section 2.22).

#### 3.6.2. Basic qualitative M_x_ method

Basic qualitative modelling was employed throughout. Results of LRs for every laboratory are listed in Supplement 6. In addition, two tests were carried out a) median value calculated across all laboratories and b) the combined data of lab 201 results (Table 7). With this method, the single contributor LR is sensitive to sample size, hence a much higher LR=191 was observed for lab 201 combined dataset.

#### 3.6.3. Comparison of continuous and M_x_ methods

*M*_*x*_ derived LRs are broadly consistent with the findings of the continuous model. To enable direct comparison (Table 7), the continuous method results are colour coded: red for ’Major’; black for ’Balanced’ and blue for ’Single’ contributor where the definition *M*_*x*_ ≥ 0.99 is applied.

1. Only ’Single’ contributors provide LRs that may be considered probative. The *M*_*x*_ method gave a result consistent with 0.5ng recovery of POI DNA for the continuous method.
2. ’Balanced’ and ’Major’ POI outcomes were similar in both methods (albeit the ’Major’ outcome gave a slightly higher LR=4).
3. With combined data (lab 201), the *M*_*x*_ method gave a higher LR=191 compared to the continuous method (LR=94 for 10ng total DNA recovery).

#### 3.6.4. Inter-laboratory variation of continuous and M_x_ method

The results from individual laboratories showed that only ’Single’ contributor DNA profiles aligned with the POI provided support for the *H*_*p*_ proposition. These results were compiled in Table 8 and ranked by LR in ascending order based on 9ng recovery.

**Table 8:** A compilation of LR results across all laboratories for experiment 2, where 0.01 - 9ng DNA are recovered (top row labels), with no background DNA. Ranked by recovery of 9ng DNA per laboratory. In the laboratory identifier column, the shaded background illustrates those labs in the middle tertial of results, Insufficient data for labs in italics

|  | DNA quantity (ng) |  |  |  |  |  |  |
| --- | --- | --- | --- | --- | --- | --- | --- |
| Laboratory | 0.01 | 1 | 3 | 4 | 5 | 7 | 9 |
| lab_21 | 6 | 8 | 6 | 6 | 6 | 5 | 5 |
| lab_6 | 9 | 17 | 13 | 12 | 11 | 10 | 9 |
| lab_13 | 8 | 14 | 18 | 19 | 19 | 20 | 19 |
| lab_17 | 13 | 19 | 24 | 25 | 26 | 26 | 26 |
| lab_18_GOF | 15 | 31 | 38 | 38 | 38 | 38 | 37 |
| lab_11 | 22 | 47 | 58 | 59 | 60 | 59 | 57 |
| lab_14 | 24 | 31 | 47 | 52 | 56 | 61 | 64 |
| lab_12 | 23 | 81 | 81 | 79 | 77 | 72 | 69 |
| lab_18_F6C | 30 | 64 | 76 | 77 | 77 | 76 | 74 |
| lab_16 | 25 | 36 | 60 | 67 | 72 | 79 | 83 |
| lab_3_ESX | 23 | 67 | 84 | 87 | 89 | 90 | 90 |
| lab_19 | 43 | 78 | 97 | 99 | 101 | 101 | 99 |
| lab_9 | 28 | 94 | 111 | 111 | 110 | 106 | 102 |
| lab_23 | 26 | 50 | 72 | 79 | 86 | 96 | 105 |
| lab_3_NGM | 23 | 64 | 90 | 96 | 102 | 110 | 116 |
| lab_22 | 54 | 73 | 117 | 136 | 154 | 186 | 215 |
| lab_4_ESI | 13 | 48 | 117 | 151 | 186 | 257 | 329 |
| lab_2_NGM | 24 | 87 | 178 | 213 | 244 | 297 | 342 |
| lab_2_ESI | 24 | 66 | 164 | 213 | 262 | 362 | 463 |
| lab_1_NGM | 26 | 137 | 279 | 341 | 402 | 517 | 629 |
| lab_4_NGM | 43 | 153 | 401 | 537 | 683 | 1001 | 1352 |
| lab_7 | 23 | 133 | 389 | 536 | 697 | 1054 | 1457 |
| lab_1_ESS | 46 | 268 | 586 | 735 | 882 | 1171 | 1459 |
| lab_8 | 27 | 139 | 437 | 606 | 785 | 1171 | 1586 |
| lab_5 | 13 | 121 | 616 | 952 | 1330 | 2188 | 3150 |
| *lab_10* | *7* | *5* |  |  |  |  |  |
| *lab_15* | *5* | *94* | *2164* |  |  |  |  |
| Medians | **24** | **66** | **90** | **96** | **101** | **101** | **102** |

The median LR across all laboratories varied from 20 - 100 according to quantity recovered, plateauing at 3ng (i.e., the LR doesn’t change with higher quantities and this was very similar to findings of combined data (lab 201) discussed in the previous section). However, there is considerable difference in LRs between laboratories: e.g., for 9ng recovery, the range is LR=5 to 3000. The observation of a ’Single’ contributor POI in the dataset (here defined as *M*_*xPOI*_ ≥ 0.99) was relatively low and found in just 41/615 samples (Pr=0.07) - i.e., the majority of results were mixtures. The individual laboratory results for the *M*_*x*_ method are listed in supplement 6 and summarised in Table 10. For experiment 2, the LR ranged between 11-173; for ’Balanced’ mixtures, results were either neutral or supported the *H*_*d*_ proposition; when the person of interest aligned with the ’Major’ LR=12 was the maximum value observed for this outcome.

For mixed samples, there is much less sensitivity to inter-laboratory variation. Results tend to either support the *H*_*d*_ proposition, or are neutral.

The following generalisations can be made:

1. LRs *>* 1, if *H*_*p*_=true are primarily achieved when a ’Single’ DNA profile is obtained that aligns with the POI.
2. The majority of results are mixtures: only 7% of samples were observed to be ’Single’ DNA profiles.
3. If the POI aligns with components in a mixture, the value of the evidence tends to be neutral or it supports the *H*_*d*_ proposition.

### 3.7. Experiment 3

#### 3.7.1. Trends for continuous and M_x_ methods

Laboratory results were combined into a single dataset ’lab 300’ and likelihood ratios calculated for both continuous and *M*_*x*_ models (table 9). Individual laboratory results are listed in Supplement 6.

**Table 9:**
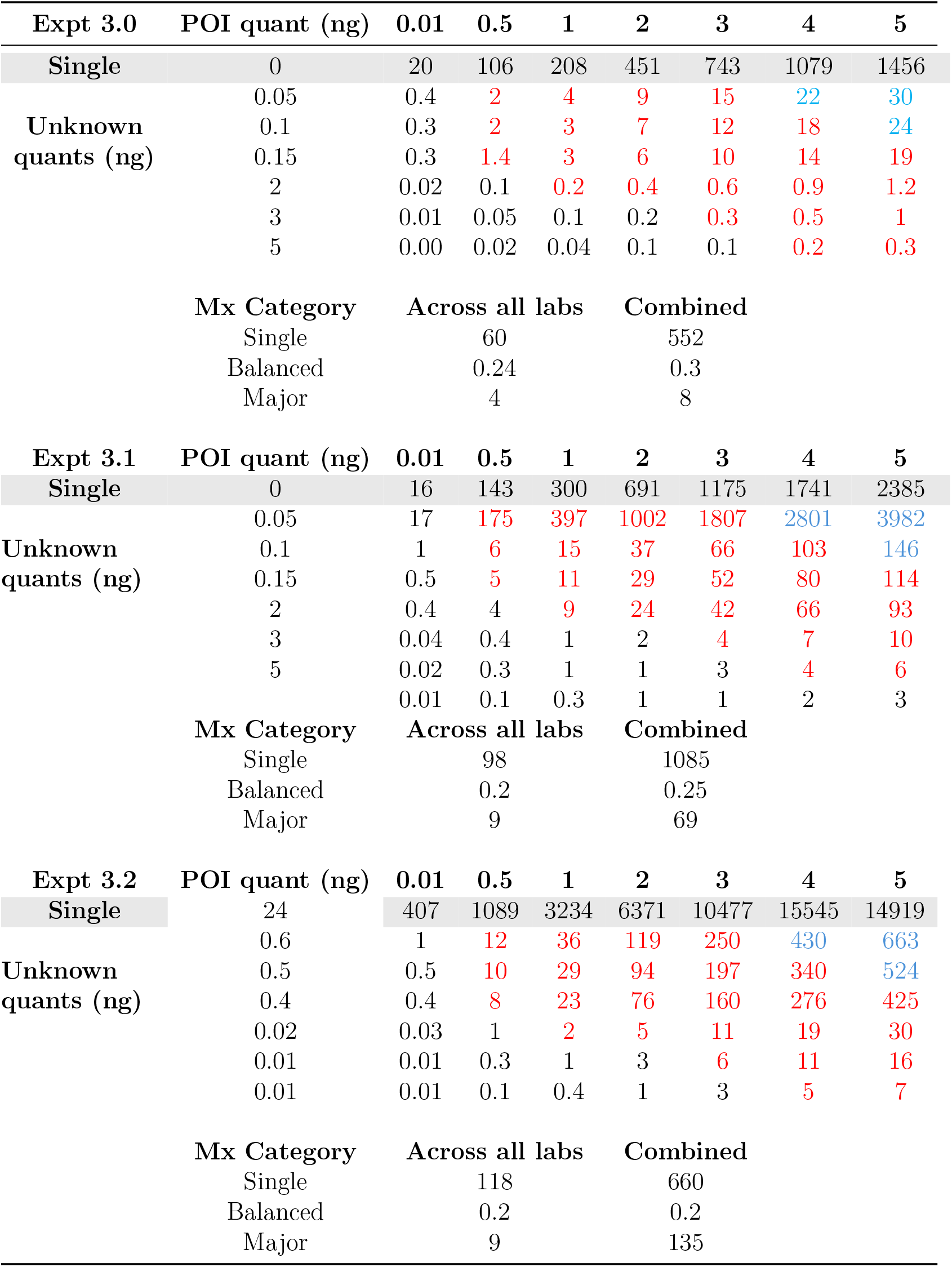
Experiment 3.0, 3.1 and 3.2 LR results from combined data ”lab 300”. Continuous model results compared to *M*_*x*_ method. For the latter, the median LR is shown ”Across all labs” along with the results from all laboratories combined (lab 300). The results between the continuous and Mx methods can be directly compared. LRs shown in black type are ”Balanced”; red type ”Major” and blue type is defined as ”Single” with the *M*_*x*_ method, but is analysed as a mixture for the continuous method

**Table 10:** Summary statistics of the *M*_*x*_ method for various categories where the POI is Single, Balanced or Major contributor. Compared with the presence/absence (P/A) binary method described in section 2.21. N=sample size

| Experiment | Statistic | Single LR | Balanced LR | Major LR | P/A Single | P/A Mixture | N |
| --- | --- | --- | --- | --- | --- | --- | --- |
| Ex_2 | median | 55 | 0.8 | 3 | 13 | 0.4 | 20 |
| Ex_2 | min | 11 | 0.3 | 1 | 3 | 0.1 | 16 |
| Ex_2 | max | 173 | 1 | 12 | 34 | 1 | 40 |
| Ex_3.0 | median | 60 | 0.2 | 4 | 18 | 0.4 | 20 |
| Ex_3.0 | min | 4 | 0.06 | 0.98 | 4 | 0.1 | 11 |
| Ex_3.0 | max | 288 | 0.8 | 18 | 85 | 1 | 35 |
| Ex_3.1 | median | 98 | 0.2 | 9 | 45 | 1 | 20 |
| Ex_3.1 | min | 3 | 0.1 | 0.5 | 2 | 0.1 | 1 |
| Ex_3.1 | max | 360 | 0.7 | 29 | 186 | 2 | 36 |
| Ex_3.2 | median | 118 | 0.2 | 9 | 49 | 0.9 | 20 |
| Ex_3.2 | min | 3 | 0.05 | 0.5 | 5 | 0.1 | 11 |
| Ex_3.2 | max | 525 | 1 | 25 | 384 | 16 | 35 |

The same trends described for experiment 2 occurred with experiment 3. LRs for ’Single’ contributors ’POI only’ are the highest, and are dependent upon the quantity of DNA recovered. In addition, the longer the time interval between the handshake and the sampling, the greater the corresponding LR value. This is not surprising, because the probability of DNA recovery of the person who shook hands quickly decreases over two hours. Compared with experiment 2, it was more likely to recover single profiles: Pr=0.18; 0.24; 0.28, respectively for zero, one, two, hours post handshake. Compared to experiment 2, if *H*_*p*_ =true, LRs are much higher for both the *M*_*x*_ and continuous models and this is driven by the lower recovery rates of FirstH (Pr secondary transfer). If the POI is a minor or aligns with components of a ’Balanced’ mixture, the evidence will tend to support the *H*_*d*_ proposition, whereas if the contributor is ’Major’ it will tend to support *H*_*p*_. LRs based upon the *M*_*x*_ method are not predicated by quantity and they tend to fall mid-range of the continuous model results (i.e., depending upon the quantity recovered, the continuous model will report LRs ranging from substantially smaller to higher values compared to the *M*_*x*_ model).

#### 3.7.2. Between laboratory variation

LR results, using the *M*_*x*_ method, for individual laboratories are summarised in Table 10. Results for the continuous model are listed in Supplement 7. There are substantial between laboratory differences for ’Single’ contributor recovery, for the *Mx* method, the *LR* ranges from 4-500, whereas for the continuous model, the variation can be three orders of magnitude for ’Single’ POI recovery and up to 2 orders of magnitude for mixtures.

### 3.8. Effect of sample size on the M_x_ method

For the *M*_*x*_ method, the highest LRs (as with the continuous method) are obtained when the POI aligns with a DNA profile from a single contributor. However, LRs are highly dependent upon the sample size. This is because many laboratories detect mixtures in all samples analysed, hence the ’Absent’ and ’Single’ categories are empty. The application of the Laplace smoothing ensures that the minimum probability of each category is defined as 1*/N* + 4 (where *N* = sample size) and this results in a lower bound *LR* = 2*N* + 7 for experiment 2; for experiment 3 the lower bound *LR* = *N* + 4. A minimum sample size *N* = 20 was specified, hence *LR*_*min*_ = 47 and 24, respectively. Increasing the sample size will correspondingly increase the minimum LR if ”Absent” and ”Single” results categories are empty (Table 12).

LRs calculated from combined data (Table 12 showed trends that paralleled results of continuous methods: For experiment 2, low (lab 600), median (lab 100) and high (lab 500) tertile laboratories showed same trends of LR differences for single contributor LRs and Major contributors - the corresponding increase in sample size has an effect on *LR*_*min*_ values. For experiment 3 (combined data for lab 300) single LRs are increased to c. LR ≈ 500 for all time intervals. If the POI was a major contributor, there is an increase from LR = 8 to LR = 73.

### 3.9. Bootstrapping data using the continuous model

The continuous method is also subject to effects of sample size, but it is not directly affected by counting events as described for the *M*_*x*_ method. To compensate for sampling uncertainty, data are bootstrapped and results are compared using the lower 5 percentile in section 3.11.

### 3.10. The importance of probabilistic genotyping software to help evaluation

Probabilistic genotyping software is used to calculate mixture proportions (*M*_*x*_). It also helps with determination of NOC. Overestimates of NOC may occur as a result of stutters. However, if exploratory analysis is undertaken and it is shown that NOC+1 has *M*_*x*_ *<* 0.01 then this is a good way to assign a functional minimum value to the parameter. Since highest LRs given activity level propositions are obtained with NOC=1, it is important to ensure that the value is not overestimated.

### 3.11. Risk analysis

Misleading evidence occurs when ground truth *Hd* = *true* scenarios give LRs *>* 1 and ground truth *Hp* = *true* scenarios give LRs *<* 1. As scientists do not give their opinion on the propositions, this phenomenon cannot be described as false positives (FP) and false negatives (FN) - Type II and Type I errors, respectively. When using LRs to provide the value of the DNA comparison, even if an LR is larger than one in a case where the DNA is from a non contributor, this cannot and should not be considered as an “error”: it is “misleading evidence” which is an inevitable consequence of the presence of uncertainty.

It is beyond the scope of this paper to carry out a detailed analysis of this kind and such a study is reserved for the future. For experiment 2, the misleading evidence for *Hp* = *true* ground truth experiments is very high because probabilities of FirstH and LastH are similar.

Here we compare two different models - a continuous model based on DNA quantity, vs a categorical model based on *M*_*x*_ values. Population genetics studies are independent of the laboratory carrying out the tests, whereas studies given activity level propositions are subject to a host of largely uncontrolled variables, all of which may have an affect upon the reported likelihood ratio. These factors include: the sampling method; the quantification method; the PCR method. Because there are so many variables, each laboratory effectively has a unique protocol. This makes comparative studies challenging. Although it is unrealistic to standardise methods, a better understanding of the effect of different methods on the LR is needed. Some standardisation may be possible (e.g., quantification method). Introduction of factors to help normalise data may be useful.

For all experiments, *LR >* 1 is notably obtained under when the profile is a ’Single’ contributor aligns with the POI (and *H*_*p*_ = *true*). For experiment 3, if the profile is multi-contributor and the ’Major’ aligns with the POI, then high LRs may be obtained if there is substantial DNA quantity recovered.

For any likelihood ratio model, the aim is to produce high LR *>* 1 when *H*_*p*_ = *true* and low LR *<* 1 when *H*_*d*_ = *true*. Ideally, calibration [17] of the likelihood ratio should be undertaken; this will help to optimise the model employed. This exploration is reserved for a future paper. As a preliminary investigation, we select all samples from the compiled data (`“Grand_compilation_v2.xlsx`) where the FirstH handler aligns either with a ’Single’ or ’Major’ contributor (Supplement 8): these conditions reflect the highest risks of wrongful attribution when *H*_*d*_ = *true*. Then, likelihood ratios were calculated using the models described earlier. The ground truth of contributor activities is known for the experiments undertaken.

For the experiment 3 series, ’Single’ contributors corresponding to the last handler were more commonly observed; they ranged between 21 − 28% for experiments 3.0 and 3.2, respectively. Because the probability of transfer of the first handler was much lower compared with experiment 2, the results provided greater support for *H*_*p*_. In addition, ’Major’ contributors corresponding to the POI, found in approximately 50% of samples, also have a strong potential to provide high LRs (table 9); conversely, ’Balanced’ contributors corresponding to the POI will provide evidence that is either neutral or supports *H*_*d*_.

### 3.12. Detailed analysis: H_d_ = true

#### 3.12.1. Experiment 2

For experiment 2, we simulated the case where the first handler has been wrongly accused. The perpetrator (always the last handler) is unknown. Recall that only ’Major’ or ’Single’ POI contributors are of interest since ’Balanced’ POI contributors give low LRs and the results tend to be neutral or support *H*_*d*_.

The distribution of contributor types is shown in table 11. For experiment 2, a total of 8% of profiles are ’Major’, and 1.4% are ’Single’. The risk of a profile supporting *H*_*d*_ = *true* is low. Just 8*/*655 samples were observed in this category. On further examination, three were low level and would not be reported. The remainder were reportable, however.

**Table 11:** Classification of findings as probabilities, separated into four categories for all experiments carried out, where the POI is LastH under Hp=True and FirstH under Hd=True.

| Hp=true |  |  |  |  |  |
| --- | --- | --- | --- | --- | --- |
| Experiment | Absent | Single | Balanced | Major | N |
| **2** | 0.05 | 0.08 | 0.45 | 0.43 | **655** |
| **3.0** | 0.05 | 0.21 | 0.26 | 0.48 | **549** |
| **3.1** | 0.07 | 0.24 | 0.14 | 0.55 | **482** |
| **3.2** | 0.06 | 0.28 | 0.10 | 0.55 | **527** |
| Hd=true |  |  |  |  |  |
| Experiment | Absent | Single | Balanced | Major | N |
| **2** | 0.14 | 0.014 | 0.76 | 0.08 | **655** |
| **3.0** | 0.29 | 0.004 | 0.67 | 0.04 | **485** |
| **3.1** | 0.51 | 0.000 | 0.49 | 0.002 | **498** |
| **3.2** | 0.54 | 0.002 | 0.45 | 0.000 | **486** |

**Table 12:** Summary statistics for the Mx method using combined laboratories.

| Experiment | Lab ID | Single LR | Balanced LR | Major LR | P/A Single | P/A Mixture | N |
| --- | --- | --- | --- | --- | --- | --- | --- |
| Ex_2 | lab_201 | 191 | 0.8 | 4 | 13 | 0.5 | 615 |
| Ex_2 | lab_100 | 287 | 0.8 | 3 | 14 | 0.5 | 273 |
| Ex_2 | lab_500 | 360 | 0.8 | 7 | 48 | 0.4 | 164 |
| Ex_2 | lab_600 | 68 | 0.9 | 4 | 7 | 0.5 | 178 |
| Ex_3.0 | lab_300 | 552 | 0.3 | 8 | 13 | 0.5 | 329 |
| Ex_3.1 | lab_300 | 1085 | 0.2 | 69 | 18 | 0.9 | 310 |
| Ex_3.2 | lab_300 | 660 | 0.2 | 135 | 41 | 1 | 301 |

Likelihood ratios given activity level propositions were calculated (Supplement 8) taking bootstraps for the 50 percentile and the 5 percentile (latter shown in parentheses in the following text). LRs were calculated relative to the combined datasets (`lab_201` and `lab_300` for experiments 2 and 3 respectively and also using the data from the originating laboratory).

Using the continuous method (experiment 2) with combined data (lab 201), all gave LRs that were less than 100(65). There was much more variation when calculated relative to the originating laboratory data, with one from lab 6 providing LR=1400, reduced to 210 when the 5 percentile boot-strap was recorded. The qualitative (originating lab) method gave low LRs. The *M*_*x*_ method LR=191 for combined labs (lab 201) and *LR <* 80 for the originating laboratory. According to a theory, known as the ”Turing expectation”, the fraction of non-contributor scenarios producing an *LR* ≥ *LR*_*x*_ is expected to be at most 1*/LR*_*x*_ [18, 19, 20]. If we compare results with the Turing expectation, out of a total of 655 observations, the expectation is that a false *H*_*d*_ = *true* test should give a maximum value of *LR* = 655. All results fulfil this requirement apart from the afore-mentioned lab 6 result that gave LR=1400. Small samples sizes have an effect on the LR and the bootstrap is a way to compensate, hence an adjustment using the 5 percentile bootstrap interval seems preferable as it reduced the LR to 210.

#### 3.12.2. Experiment 3

For experiment 3, the experimental design differs in that the defence proposition asserts that the last handler shook hands with the defendant, and his/her DNA was secondarily transferred. Compared to experiment 2, the probability of recovery of a major or single contributor, by secondary transfer is much lower (Table 11). Consequently, LRs are much higher for both ’Single’ and for ’Major’ profiles where *H*_*p*_ = *true* (Table 9)

For experiment 3.0, there were 17 samples where first handler was the ’Major’ contributor and two which were ’Single’ contributors. However, the latter can be eliminated because they were very low level and would not be reported when considering sub-source propositions. This leaves ’Major’ contributors to consider. For the continuous method, likelihood ratios were generally less than one or neutral for both the combined lab 300 data as well as originating lab calculations. Taking the 5 percentile bootstrap reduced highest observations from lab 2 from LR=17 to 0.6 and LR=24 to 2.5 respectively. Although the *M*_*x*_ Method also gave low LRs for the ’at risk’ dataset, compared to the continuous method, they were always higher *LR >* 1, LR=8 for combined data (lab 300) and ranged between LR=1-6 for the originating lab analysis (Supplement 8). These LRs are too low to be considered discriminative for reporting purposes, the inability to take account of quantity reduces the efficacy of the method.

Turning to experiment 3.1 there was only one observation of a major profile which gave a low LR with the continuous method LR=4.6(1.8) `lab_300`, and LR=49 with the *M*_*x*_ method. For experiment 3.2 a single contributor was recovered from lab 10 which gave a high LR as a result. However, the profile was low level and would not be reported given sub-source propositions. In addition it was noted that the contributor recovered was classed as a very high shedder because of a skin ailment. Shedder status was not taken into account in this study, but it is probably important to do so in future work.

To summarise, it is very unlikely to recover DNA that gives high LR from FirstH in experiment 3, especially if more than one hour has elapsed since contact and sampling.

### 3.13. H_p_ = true analysis

A small selection of results based on experiment 1 were taken. This simulated the situation where *H*_*p*_ = *true* and *H*_*d*_ = *false*. The samples chosen are all ’Major’ or ’Single’ contributors (Supplement 9). A comparison was made between lab 3 ESX, and combined data from lab 201 and 300 compilations.

#### 3.13.1. Experiment 2

Results in support of *H*_*p*_ were only obtained when the recovered DNA appeared to be from single contributor and that it aligned with the POI. In terms of magnitude of the *LR*, there was little difference between continuous and *M*_*x*_ methods. For lab 3 ESX results (as a representative median performing laboratory) - continuous 50 percentile: LR=43-97; *M*_*x*_ method: LR=70, otherwise major profiles all gave results that were neutral (*LR* ≈ 1). For lab 300 results, *LR*s were again similar, but the main difference was for the *M*_*x*_ method where major profiles all provided *LR* = 4 and single contributors *LR* = 191. The *M*_*x*_ LR method is sensitive to sample size and does not take DNA quantity into account.

#### 3.13.2. Experiment 3

Compared to experiment 2, *LR*s are at least an order of magnitude greater for experiment 3.0 and two orders of magnitude for experiment 3.1. There were too few data for experiment 3.2 to carry out an analysis; this was only feasible when data were combined to make a large data-set in lab 300. In lab 3 ESX, single contributors can reach LR=10,000 (750) for the continuous method, corresponding to LR=176 for the *M*_*x*_ method (experiment 3.1). For lab 300, the *M*_*x*_ method gave single contributor LRs up to LR=1000; the major contributor *LR* ≈ 135 for experiment 3.2 was substantially greater than for lab 3 ESX,again reflecting the effect of increased sample size on this method. The 5 percentile results of the continuous method were a closer fit to the *M*_*x*_ method.

### 3.14. A closer look at the probability of transfer, persistence and recovery

Previous workers (including the authors of this paper) have previously treated the ’transfer, persistence, recovery’ (TPR) parameter, as an all-encompassing probability such that:

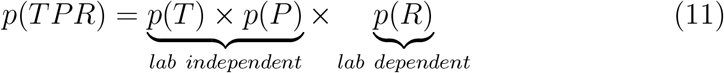

If *p*(*TPR*) has been assigned by laboratory A, an assumption may be made that it can be adopted by laboratory B, without any further work or validation. Indeed, the work of laboratory A may be cited in court by laboratory B as a reason to support some outcome.

There is an expectation that probabilities of transfer and persistence are *independent* of the laboratory and therefore should be applicable across laboratories for a given evidence type. However, the waters are muddied by the probability of recovery, which is a laboratory *dependent* parameter. Furthermore, the variation in recovery is considerable - quantity varies between 200pg and 5ng (a range of 2,500%).

Consequently, we conclude that future work must address the issue of laboratory dependent recovery. The difficulty is that to date there are no experimental designs described in the literature. This will consequently be a focus of the ReAct II project: where the intention is to determine a recovery factor using a simple experimental design that can be easily followed by other laboratories.

### 3.15. Generalisations that can be made with regard to the findings

Where single contributors were recovered (*Hp* = *true*), corresponding LRs increased progressively: experiment 2 *<* experiment 3.0 *<* experiment 3.1 *<* experiment 3.2; driven by progressive reduction in probability of FirstH (either first handler for experiment 2 or secondary transfer for experiment 3). There were considerable differences in LRs between laboratories of up to two orders of magnitude for the continuous model (dependent on recovery quantity), and less for the qualitative *M*_*x*_ method (independent of recovery quantity). The lower LR laboratories either had lower DNA recovery, and therefore fewer data, or *Pr*(FirstH) was closer to *Pr*(LastH); converging mixture proportions in experiment 2 resulted in reduced *LR*s. Each laboratory uses a different method to recover DNA; some are more efficient, and this certainly has an effect upon probability of recovery, mixture proportions and the likelihood ratio. This interaction will be explored in a subsequent paper.

When ’Balanced’ results are analysed, defined as *POI < Mx* = 0.7, LRs were usually ≤ 1 for all experiments. This is an important finding that we can generalise (i.e., it is a valid observation that can be applied to all laboratories).

If the POI aligns with a balanced mixture or with a minor contributor (*Mx <* 0.07) then the evidence does not support the prosecution case. If the POI aligns with the major, then the value of the evidence depends upon the quantity of DNA recovered. For experiment 2, the evidence is either neutral, or supports *H*_*d*_. For experiment 3, the evidence usually supports *H*_*p*_, but the LR is very dependent upon the quantities recovered. This can only be properly assessed using the continuous model.

The *M*_*x*_ method does not take into account the DNA quantity (Table 9); the *M*_*x*_ method is sensitive to sample size, hence LRs are considerably increased when data are combined (lab 300). Compared to the continuous method, the *M*_*x*_ method may overstate the value of the evidence when low quantities of DNA are recovered, and understate when high quantities are recovered.

Similarly, the presence of ’Major’ POI contributor for experiment 2 has little value in support of the *H*_*p*_ proposition. For experiment 3, the support is closely associated with the quantity of DNA recovered. Low quantities provide low support for *H*_*p*_ whereas higher quantities give much higher LRs (table 9). The lower recovery of secondarily transferred DNA (FirstH) in one and two hour post-handshake experiments 3.1, 3.2 resulted in correspondingly higher LR values.

### 3.16. On the use of proxy experiments

In casework, the position of the defence is not necessarily forthcoming; in fact this was the usual position after reviewing case examples from participant laboratories in Supplements 2,3. Indeed, the defence are under no obligation whatsoever to provide an alternative proposition. The defendant is considered to be innocent and the burden of proof is entirely the responsibility of the prosecution. Consequently, the practitioner must be extremely cautious when putting forward propositions on behalf of the defendant. A good example is provided by experiment 2, where it is accepted that the defendant owns the tool, thereby providing an agreed method of DNA transfer - the dispute between *H*_*p*_ and *H*_*d*_ is narrowed to the question of whether the defendant was the first or the last handler of the tool. The framework is complete.

Experiment 3 is more complex. Case circumstances typically encountered in casework (Supplements 2,3) are: the defendant denies ownership of the tool and has no legal obligation or information to help the scientist form a view about how the DNA may have transferred to the item. The legal framework requires that the prosecution prove the case beyond reasonable doubt. There is no reverse burden of proof on the defence to prove otherwise. Whereas the prosecution proposition is usually straight forward: the defendant forced the door with the tool; the alternative defence proposition is that he/she had nothing to do with the offence - he/she never held the tool. However, if the defendant’s DNA is present (and agreed by both parties), there is often no information to help the court with the question of how transfer occurred.

At this point there is an option for scientist to state that he/she is unable to help the court further, other than to list all possible options. However, this is unsatisfactory, since the court is left to its own devices to make the ultimate decision of guilt/innocence on the basis of a DNA profile comparison, where it is agreed is that the DNA comes from the defendant. Therein lies the greatest danger in the propagation of miscarriages of justice. The scientist is obligated to help the court. A possible way forward is to consider the use of proxy experiments.

#### 3.16.1. Design of proxy experiments

It must be stressed that when scientists give evidence to courts they should not directly address issues of ”transfer”; instead, propositions should describe the alleged activities, for example ”the defendant shook hands with an individual” or ”the defendant had some form of social activity”. The probabilities of different modes of transfer events are conditioned on activities and this is illustrated by implementation in Bayesian Networks (i.e., we discuss ”transfer” only in relation to probabilities used to inform the Bayesian network which are conditioned by the activity nodes). The distinction is important. In court, the discussion is limited to the value of the findings given propositions outlined above. As a general guideline or test, propositions should ideally be set before the findings are known.

As stated earlier, for experiment 3, the given prosecution’s view was that there had been direct transfer to the tool. The given defence view was that some kind of indirect transfer took place and there was no direct transfer. The method of indirect transfer was unknown: it could be secondary transfer, tertiary or higher order of transfer events.

An alternative strategy is to maximise the probability of the evidence, given the defence proposition, by carrying out a reasonable experiment, designed to maximise the transfer of DNA under *H*_*d*_. This is achieved by hand-shaking experiment 3.0. This method demonstrably transfers more DNA compared to higher order tertiary transfer and will therefore be conservative - the defence view is: ”the defendant did not handle the tool; it is not known who handled the tool”; it is not known how or when the defendant’s DNA was propagated. The results of experiments 3.1 and 3.2 are also relevant to the debate. Under this scheme, there is no need for the defence to engage in discussions about *specific* methods of DNA transfer. Whether there was an *opportunity* is solely a matter for court consideration.

The function of the scientist is to make sure that the court is given all the information relative to the DNA analysis and its meaning, as this is necessary to make a decision. It is not the function of the scientist to filter information based on preconceived perceptions of relevance. This is known as *confirmation bias*. The court, as the fact finder, decides *how* to use the information and the weight to place upon it.

### 3.17. Standardisation

Both the continuous method, and the *M*_*x*_ method require analysis using probabilistic genotyping (PG) tools such as EuroForMix. Other tools such as STRmix could also be used for this purpose. In order to facilitate reporting given activity level propositions, laboratories will need to implement PG methods.

When large amounts of data are analysed it is essential that as much of the process is automated as possible. This was achieved by asking laboratories to prepare concatenated data in a standard format that could be analysed by EuroForMix. The ShinyRFU() app was then used to automate analysis, outputing *M*_*x*_ values and other data. Laboratories also submitted spreadsheets with details of DNA quantities, numbers of contributors etc, again in a standard format that facilitated automated analysis.

### 3.18. Difficulties of organising large collaborative exercises

The main problem encountered is that we did not anticipate the huge variation in results that were obtained. This obviated the original aim to tie the RFU results with quantification method as described by [11]. It was also found that laboratories that did not attend a meeting to discuss the protocols were more likely to make mistakes - data from one laboratory were removed because the wrong protocol was followed. Consequently, a lot of effort was expended in resolving data errors. For example, some labs reversed first handler / second handler notation. These errors are fairly easy to spot, but of course nothing was changed without the originating laboratory confirmation.

It is likely that some errors remain in the dataset. If detected, the data may be altered. If changes are made, to maintain continuity, databases will be updated with version numbers. In addition, we will add data from other laboratories interested in contributing. The project will continue to add new data and evolve.

## 4. Conclusion

### 4.1. Reporting LRs given activity level propositions: Future plans

To return to the issue of whether laboratory A can use data of laboratory B for evaluation purposes (calculation of likelihood ratios), this project lays the groundwork for future work. There needs to be much greater attention to the probability of recovery parameter because it is laboratory dependent and the range is considerable. The ReAct II project will provide a focus on the parameter to identify a way forward. Once this has been achieved, the data can be revisited. In the interim, a possible way forward is to use the combined datasets in order to derive likelihood ratios. These data provide an ’average’ expectation of results. In addition, to help accommodate small datasets, it is preferable to report results using 5 percentile bootstraps for the continuous method. However, these are suggestions rather than firm recommendations. A better way forward is to implement full Bayesian models that properly take account of informative prior distributions (e.g. using beta binomial models outlined in section 2.16).

To summarise, there is a particular challenge to reconcile the huge differences in laboratory techniques and interpretation methods so that evaluation given activity level propositions takes this variation into account. This will be a focus of future publications.

The findings of this work can be summarised as follows.

### 4.2. Experiment 2

From basic qualitative modelling: if an individual handles a screwdriver, then there is approximately 95% probability of DNA transfer. If two individuals handle a screwdriver, then the probability of recovery of the first handler drops to 88%. In approximately 43% of cases, the last handler was ’Major’, compared to 8% of cases for the first handler. Likelihood ratios for ’Major’ contributors are low. However, results allow discrimination when the DNA profile of the POI aligns with the DNA profile of a trace assigned as from a single contributor (found in 8% of cases when *H*_*p*_ is true). There were six examples of experiments where the first handler was the single contributor, and consequently high LRs were also obtained in these cases. Although the results were well within Turing expectations, calibration of models is desirable to ensure that results are not misleading.

### 4.3. Experiment 3

Secondary transfer was observed in c.69% of cases provided that the screwdriver was held immediately after handshake, otherwise it drops to c.48% percent after one hour post handshake. Consequently higher LRs were obtained provided that the recovered contributor was ’Major’ or ’Single’ (about 70% of cases). Incidences of misleading LRs (LR *>* 1) were restricted to single examples in experiments 3.1 and 3.2 where the former was a major profile with low LR given activity level propositions, and the later a single profile, but of too low quality to report given sub-source level propositions.

Consequently, if a ’Major’ or ’Single’ profile aligning with a person of interest is recovered from a crime scene, where the defence proposition is that some form of social contact has occurred at least one hour previously, the DNA results will usually support the proposition that the person has handled the tool (the recovered DNA always aligned with the last handler of the screwdriver).

### 4.4. What about the LR?

As a final comment, it is worth noting that the theory is very well developed but the application lags somewhat behind. We have highlighted an issue with reproducibility of experiments. Unfortunately, the generalisations listed above do not translate into likelihood ratios, given activity level propositions, which can be adopted by laboratories that have not undertaken experimentation. There is no short-cut at present, but this is the subject of further research currently being undertaken by our laboratories.

## Acknowledgements

Funding for this Collaborative Project was received from the European Union’s Internal Security Fund -Police (ISFP) – with Grant Agreement title: Competency, Education, Research, Testing, Accreditation, and Innovation in Forensic Science” [CERTAIN-FORS] ISFP-2020-AG-IBA-ENFSI and number: 101051099. We acknowledge support from ENFSI and the ENFSI DNA Working Group Chairman Sander Kneppers. The authors thank the Forensic Genetics and Casework section of the Institute of Legal Medicine, Medical University of Innsbruck, for excellent technical assistance.

## Footnotes

1 The interested user is free to use any criteria to decide whether a contributor is present or absent: accordingly, the spreadsheet is designed so that the data are easily filtered.

## References

[1] B. Kokshoorn, L. H. Aarts, R. Ansell, E. Connolly, W. Drotz, A. D. Kloosterman, L. G. McKenna, B. Szkuta, R. A. van Oorschot, Sharing data on DNA transfer, persistence, prevalence and recovery: arguments for harmonization and standardization, Forensic Science International: Genetics 37 (2018) 260–269.

[2] A. Gosch, C. Courts, On DNA transfer: the lack and difficulty of systematic research and how to do it better, Forensic science international: Genetics 40 (2019) 24–36.

[3] R. A. Van Oorschot, K. N. Ballantyne, R. J. Mitchell, Forensic trace DNA: a review, Investigative genetics 1 (2010) 1–17.

[4] R. A. Van Oorschot, B. Szkuta, G. E. Meakin, B. Kokshoorn, M. Goray, DNA transfer in forensic science: a review, Forensic Science International: Genetics 38 (2019) 140–166.

[5] K. Steensma, R. Ansell, L. Clarisse, E. Connolly, A. D. Kloosterman, L. G. McKenna, R. A. van Oorschot, B. Szkuta, B. Kokshoorn, An inter-laboratory comparison study on transfer, persistence and recovery of DNA from cable ties, Forensic Science International: Genetics 31 (2017) 95–104.

[6] B. Szkuta, R. Ansell, L. Boiso, E. Connolly, A. D. Kloosterman, B. Kokshoorn, L. G. McKenna, K. Steensma, R. A. van Oorschot, Assessment of the transfer, persistence, prevalence and recovery of DNA traces from clothing: an inter-laboratory study on worn upper garments, Forensic Science International: Genetics 42 (2019) 56–68.

[7] B. Szkuta, R. Ansell, L. Boiso, E. Connolly, A. D. Kloosterman, B. Kokshoorn, L. G. McKenna, K. Steensma, R. A. van Oorschot, DNA transfer to worn upper garments during different activities and contacts: An inter-laboratory study, Forensic Science International: Genetics 46 (2020) 102268.

[8] N. Leuenberger, N. Jan, T. Kuuranne, V. Castella, Characterization of DNA concentration in urine and dried blood samples to detect the c. 577 deletion within the EPO gene, Drug Testing and Analysis (2024).

[9] Ø. Bleka, G. Storvik, P. Gill, EuroForMix: An open source software based on a continuous model to evaluate STR DNA profiles from a mixture of contributors with artefacts, Forensic Science International: Genetics 21 (2016) 35–44.

[10] P. Gill, Ø. Bleka, O. Hansson, C. Benschop, H. Haned, Forensic practitioner’s guide to the interpretation of complex DNA profiles, Academic Press, 2020.

[11] P. Gill, Ø. Bleka, A. E. Fonneløp, Limitations of qPCR to estimate DNA quantity: An RFU method to facilitate inter-laboratory comparisons for activity level, and general applicability, Forensic Science International: Genetics 61 (2022) 102777.

[12] J. W. Tukey, Exploratory Data Analysis, Addison-Wesley, 1977.

[13] P. Gill, T. Hicks, J. M. Butler, E. Connolly, L. Gusmão, B. Kokshoorn, N. Morling, R. A. van Oorschot, W. Parson, M. Prinz, et al., DNA commission of the International society for forensic genetics: Assessing the value of forensic biological evidence-Guidelines highlighting the importance of propositions. Part II: Evaluation of biological traces considering activity level propositions, Forensic Science International: Genetics 44 (2020) 102186.

[14] L. Carrara, T. Hicks, L. Samie, F. Taroni, V. Castella, DNA transfer when using gloves in burglary simulations, Forensic Science International: Genetics 63 (2023) 102823.

[15] H. Johannessen, P. Gill, G. Shanthan, A. E. Fonneløp, Transfer, persistence and recovery of DNA and mRNA vaginal mucosa markers after intimate and social contact with Bayesian network analysis for activity level reporting, Forensic Science International: Genetics 60 (2022) 102750.

[16] C. C. Benschop, M. Slagter, J. H. Nagel, P. Hovers, S. Tuinman, F. E. Duijs, L. J. Grol, M. Jegers, A. Berghout, A.-W. Van der Zwan, et al., Development and validation of a fast and automated DNA identification line, Forensic Science International: Genetics 60 (2022) 102738.

[17] D. Ramos, J. Gonzalez-Rodriguez, Reliable support: Measuring calibration of likelihood ratios, Forensic science international 230 (1-3) (2013) 156–169.

[18] D. Taylor, J. Buckleton, I. Evett, Testing likelihood ratios produced from complex DNA profiles, Forensic Science International: Genetics 16 (2015) 165–171.

[19] P. Gill, J. Curran, C. Neumann, A. Kirkham, T. Clayton, J. Whitaker, J. Lambert, Interpretation of complex DNA profiles using empirical models and a method to measure their robustness, Forensic Science International: Genetics 2 (2) (2008) 91–103.

[20] P. Gill, Ø. Bleka, T. Egeland, Does an English appeal court ruling increase the risks of miscarriages of justice when complex DNA profiles are searched against the national DNA database?, Forensic Science International: Genetics 13 (2014) 167–175.

