## Supplement 1 for "The ReAct project: Analysis of data from 23 different laboratories to characterise DNA recovery given two sets of activity level propositions"

**Supplementary material 1**

List of laboratory combinations

1) Lab 201 Experiment 2: All laboratories were combined for this category except for labs 10 and 15, because of insufficient data
2) Lab 500 Experiment 2 Top tertile of LR results:
   i. "lab_2_ESI",
   ii. "lab_1_ESS",
   iii. "lab_1_NGM",
   iv. "lab_4_NGM",
   v. "lab_7",
   vi. "lab_8",
   vii. "lab_5"
3) Lab 100 Experiment 2 mid tertile of LR results
   i. "lab_2_NGM",
   ii. "lab_3_ESX",
   iii. "lab_3_NGM",
   iv. "lab_16",
   v. "lab_22",
   vi. "lab_23",
   vii. "lab_9",
   viii. "lab_19",
   ix. "lab_18_F6C",
   x. "lab_4_ESI"
4) Lab 600 Experiment 2 low tertile of LR results
i. "lab_21",
ii. "lab_6",
iii. "lab_13",
iv. "lab_17",
v. "lab_18_GOF",
vi. "lab_14",
vii. "lab_11",
viii. "lab_12"
5) Lab 300 Experiment 3
   i. "lab_9",
   ii. "lab_21",
   iii. "lab_15",
   iv. "lab_17",
   v. "lab_13",
   vi. "lab_10",
   vii. "lab_18_F6C",
   viii. "lab_6",
   ix. "lab_14",
   x. "lab_3_ESX",
   xi. "lab_8",
   xii. "lab_12",
   xiii. "lab_7",
   xiv. "lab_3_NGM",
   xv. "lab_2_NGM",
   xvi. "lab_4_NGM"
