## Supplement 2 for "The ReAct project: Analysis of data from 23 different laboratories to characterise DNA recovery given two sets of activity level propositions"

**ReACT compilation of cases**

The purpose of the compilation was to give an overview of the kinds of cases that are reported in laboratories where a tool is used in a burglary. There were 26 examples provided (reproduced below). Some labs submitted multiple examples. The majority of labs reported cases where a varieties of tools were used and recovered at the crime scene. Types of tools included screwdrivers, crowbars, a stone. There were also reports involving guns and a hand grenade. Some cases involve theft of gloves which were subsequently use in the offence.

Sub-source propositions always indicate a defendant (POI) as the donor of DNA

Prosecution propositions always allege that the defendant used the tool in the course of the robbery.

Defence propositions vary, but have been subdivided into two broad categories;

a) Type 1 case (direct transfer): The defendant accepts ownership of the tool but claims that it was lost or stolen some time previously, and the offence was carried out by an unknown person. Or in one case the defendant suggests that he may have touched the tool previously, but denies the offence
b) Type 2 case (indirect transfer): The defendant denies both ownership and handling of the tool. Usually there is no alternative explanation put forward. However, it is logical that indirect transfer can be the only defence alternative, especially when there is no link between the defendant and the victim, or the premises where he/she lives.

In total, there eight type 1 cases, eighteen type 2 cases reported, and two indeterminate cases where both type 1 and type 2 propositions are formulated.

See spreadsheet (Supplement 3) for compilation (originating labs not disclosed here).

**Example 1**

1) What are the allegations made by the prosecution and the defence?

Mr X was responsible for the burglary.

Mr X was not involved in the burglary, stating that he had lost, or had stolen from him, countless items of clothing, including gloves.

2) What is the uncontested information?

A wooden shed in the garden of the address was broken into, a tool taken from within and used to gain entry to the house. A search was made of a number of rooms in the house. The occupant awoke and disturbed the individual. A number of personal items were stolen. A glove and a scarf that did not belong to the occupant were recovered from the property and a garden tool was found just outside the property. The tool was believed to have been used to gain entry.

The police officer who seized the items did so as follows: Glove seized at 00:30, scarf seized twenty minutes later at 00:50 moments before the tool, also seized at 00:50. The officer was wearing gloves when he seized the items, but did not change his gloves between the seizure of these items. He could not recall where he touched the tool in order to seize it. These items were all seized within approximately an hour of the incident.

3) What is the Contested information?

Mr X denies that he was involved in the burglary and does not accept that he was present or that he participated in any way. He was homeless at the time of the incident and kept his possessions in a bag. He had lost or had stolen from him countless items of clothing, including gloves, due to the manner in which he lived.

4) Results

**Glove**: High level clear and complete major male profile matching Mr X obtained from a sample of possible cellular material from the inside surface. Routine statistical evaluation conducted, likelihood ratio in excess of one billion. Contribution of DNA from at least one further individual at much lower level. Suitable for comparisons if further reference DNA profiles supplied.

**Scarf**: High level clear and complete major male profile matching Mr X obtained from a sample of possible cellular material from one side of the scarf. Routine statistical evaluation conducted, likelihood ratio in excess of one billion. Contribution of DNA from at least two further individuals at much lower level. Suitable for comparisons if further reference DNA profiles supplied.

**Tool**: Low level mixed result, indicating the presence of DNA from at least two individuals, obtained from a sample taken from the handle. No clear major contributor identified. Majority of the components detected matched the corresponding components in the profile of Mr X such that he could be considered as a potential contributor of DNA to the result. Suitable for statistical evaluation using probabilistic genotyping software, but not conducted. Trace amount of DNA not attributable to Mr X also present.

5) Assumptions

For the purposes of the activity level evaluation, it was assumed that the DNA matching Mr X on all three items had indeed originated from him.

6) The propositions:

Mr X wore the glove and the scarf and handled the tool at the time of the burglary, as alleged.

Mr X was not involved in the burglary and had lost or had stolen from him countless items of clothing, including gloves.

**Example 2**

What are the allegations made by the prosecution and the defence?

Mr X was responsible for the burglary.

Mr X was not involved in the burglary, stating that he never seen or handled the tools in question. He had a Stanley knife, but did not know where it was now.

1) What is the uncontested information?

Licenced premises, The Parkers Arms, was broken into. Entry was gained by the removal of single pane windows and wooden beading. CCTV footage showed the offender carrying a pair of bolt cutters and not wearing gloves. The bolt cutters were used to attempt to attempt to break the safe – this was unsuccessful. The bolt cutters were found outside in the pub beer garden the same day along with a pair of garden shears and half a Stanley knife casing.

Mr X lived in the residential flat above The Parkers Arms. The other half of the Stanley knife casing was found under his bed.

2) What is the Contested information?

Mr X denies that he was involved in the burglary. He was shown photographs of the tools seized (including the bolt cutters) and denied ever seeing them or handling them. He said that he has a Stanley knife, but did not know where it was now. He could not answer why half of a Stanley blade (casing?) was in his room and the other half was outside.

3) Results

**Bolt cutters**: A mixed result, indicating the presence of DNA from at least five individuals, including at least one male, was obtained from a sample taken from the handles of the bolt cutters. Profile of Mr X fully represented such that he could be a possible contributor of DNA to the result. Result statistically evaluated with probabilistic genotyping software (STRmix) and LR of in excess of one billion times more likely if Mr X contributed rather than he did not obtained (using Mr X and four unknowns / five unknowns).

4) Assumptions

For the purposes of the activity level evaluation, it was assumed that the DNA matching Mr X had indeed originated from him and that there was a chance (however small) that some DNA attributable to him could have transferred indirectly to the bolt cutters given the close proximity of his home address and that the item was recovered from a public area, next to part of a Stanley knife which is believed to belong to him.

5) The propositions:

Mr X has handled the bolt cutters.

Mr X has not handled the bolt cutters.

**Example 3**

1) What are the allegations made by the prosecution and the defence?

Mr and Mrs X were responsible for the burglaries.

Mr and Mrs X were at the location in order to meet a male regarding a potential job opportunity for Mr X. The items that were in their car that were believed to have been stolen were there in order to be given to said male.

2) What is the uncontested information?

Series of burglaries occurred at business properties overnight. Entry was gained to all three properties with force and tool marks were found on all of the doors. Swabs taken from a Dewalt tool recovered from one of the properties and swabs taken from the handle of a knife found by the point of entry at a second property. Swabs taken approximately six and a half hours after the alleged incidents. Mr and Mrs X found outside a shop nearby where an alarm was sounding. Tools found on the person of Mr X. Stolen items found at the home address of Mr and Mrs X.

3) What is the Contested information?

Mr and Mrs X state that they were there at that time in the morning (4am) as they were due to meet a male regarding a job opportunity for Mr X. The stolen items located in their vehicle were there to be given to said male. Mr X ‘finds things and picks them up’ and this is what happened with the tools in a black bag. Mr X stated to officers initially that he had tools on his person as ‘I steal’, but in interview he stated that he did not steal or burgle.

4) Results

**Swabs from knife handle**: A low level mixed result, indicating the presence of DNA from at least three individuals, including at least one male, was obtained. Mr X fully represented such that he could not be excluded as a possible contributor. Result statistically evaluated with probabilistic genotyping software (STRmix) and LR of approximately one hundred million times more likely if Mr X contributed rather than he did not obtained (using Mr X and two unknowns / three unknowns). Not possible to reliably determine whether or not Mrs X contributed.

**Swabs from Dewalt tool**: A low level mixed result, indicating the presence of DNA from at least three individuals, including at least one male, was obtained. A number of the components in the profile of Mr X represented such that he could not be excluded as a possible contributor. Result statistically evaluated with probabilistic genotyping software (STRmix) and LR of in excess of one billion times more likely if Mr X contributed rather than he did not obtained (using Mr X and two unknowns / three unknowns). Not possible to reliably determine whether or not Mrs X contributed.

5) Assumptions

For the purposes of the activity level evaluation, it was assumed that the DNA matching Mr X had indeed originated from him and that there was a chance (however small) that some DNA attributable to him could have transferred indirectly to the items given the relatively close proximity of where Mr X was when he was arrested and the number of unknowns in his defence (e.g. when he was last in the area, if at all, and whether he had ever been inside these premises before). It was assumed, for the purposes of the evaluation, that Mr X had not entered the premises from which the items were recovered.

6) The propositions:

Mr X had contact with the knife and the Dewalt tool.

Mr X did not have contact with the knife or the Dewalt tool and had not entered the relevant premises, but had been waiting for someone nearby.

**Example 4**

7) What are the allegations made by the prosecution and the defence?

- Mr X used a bolt cutter to cut the lock of a storehouse from which items were stolen between 15/01/2020 (18:30)-16/01/2020 (06:55).
- Mr X states that he has never seen the bolt cutter before and claims that someone who has used his gloves deposited his DNA onto the bolt cutter.

8) What is the uncontested information?

According to the information provided by the police investigator, the lock of the door of a storehouse was cut-open using a bolt cutter and items were stolen from the storehouse. The theft took place between 15/01/2020 (18:30)-16/01/2020 (06:55). At 06:55, 16/01/2020, the owner of the storehouse reported the theft to the police. Upon arrival at the crime scene on 16/01/2020, the police investigators at 07:50 found a bolt cutter near the storehouse. The owner stated that the bolt cutter did not belong to him and he has never seen it before. The bolt cutter was packaged and sent for DNA analysis. There was no CCTV camera in the area, therefore, no information could be provided regarding the movements of the perpetrator(s) and whether he wore gloves or not.

9) What is the Contested information?

According to Mr X, he has nothing to do with the bolt cutter and the theft of the storehouse. He states that he never owned a bolt cutter and he claims that his DNA was transferred onto the bolt cutter by an unknown individual who wore his used gloves.

10) Results

DNA extracted from the handles of the bolt cutter was amplified using the PowerPlex-ESX17 STR kit. Two separate amplifications were carried out. In one of the amplifications the alleles of the SE33 locus were below the set threshold of 100 RFUs (ABI-3130xl). A mixed DNA profile was obtained from each of the two amplifications of the aforementioned recovered DNA. Mr. X matched the major profile. There were two and one alleles that did not belong to Mr.X in the mixed DNA profile from the first and second amplification, respectively.

11) Assumptions

- Based on the results described in the DNA report, the DNA profile of Mr. X. is compatible with the major profile for 15 of 16 loci of the first amplification and 16 of 16 loci of the second amplification. The Amelogenin locus indicating X-Y genotype in both amplifications is excluded from the above counting of loci. The source of the DNA is not contested. Therefore, it is considered that the major proportion of the DNA recovered from the seized bolt cutter is from Mr. X. There is also a very small amount of DNA from an unknown contributor present.
- The lock of the door was cut with a bolt cutter by a person who may or may not have worn gloves.
- No specific information on Mr X shedder status exists, it was considered that he or an alternative offender are compatible with respect to shedder status.

12) The propositions:

Set-1

- Mr X, while **not** wearing gloves, cut the lock of the door of the storehouse with a bolt cutter between 15/01/2020 (18:30) and 16/01/2020 (06:55).
- An unknown person, while wearing the gloves of Mr.X, cut the lock of the door of the storehouse with a bolt cutter between 15/01/2020 (18:30) and 16/01/2020 (06:55).

Set-2

- Mr X, while wearing gloves, cut the lock of the door of the storehouse with a bolt cutter between 15/01/2020 (18:30) and 16/01/2020 (06:55).
- An unknown person, while wearing the gloves of Mr.X, cut the lock of the door of the storehouse with a bolt cutter between 15/01/2020 (18:30) and 16/01/2020 (06:55).

**Example 5**

13) What are the allegations made by the prosecution and the defence?

Burglary at a residential property and mark is swabbed from the window frame at the point of entry and tools are left inside that do not belong to the owner; a safe had been broken into and property and cash taken. The swab of the mark and the handle of a chisel were sampled and analysed.

- Suspect used tools to break in and was in contact with window at point of entry as he gained entry to the property
- Suspect denied entry and stated that someone had stolen his tools from his shed 6 weeks prior to the burglary

14) What is the uncontested information?

It is accepted that the DNA recovered from the contact mark on the window and on the handle of the tool originates from the suspect and that the tools originally belonged to the defendant.

It is agreed that whilst no blood was found, it is possible that some other body fluid such as saliva might be present (this was not tested for) – therefore cannot say from what biological source the DNA has originated from.

Only one tool handle analysed and it is agreed that it is not known how much DNA from suspect on the untested tools.

Do not know what made the mark i.e. whether contact with skin or fabric.

15) What is the Contested information?

Suspect states that someone else had stolen his tools and used them in the commission of the burglary. The implication is that the tools have transferred his DNA to the window frame (secondary) or that the DNA transferred from the tool, to the hands of the offender and from there to the window frame.

16) Results

A complete major profile with minor components commensurate with background was obtained from the contact marks which matches suspect (Billion times)

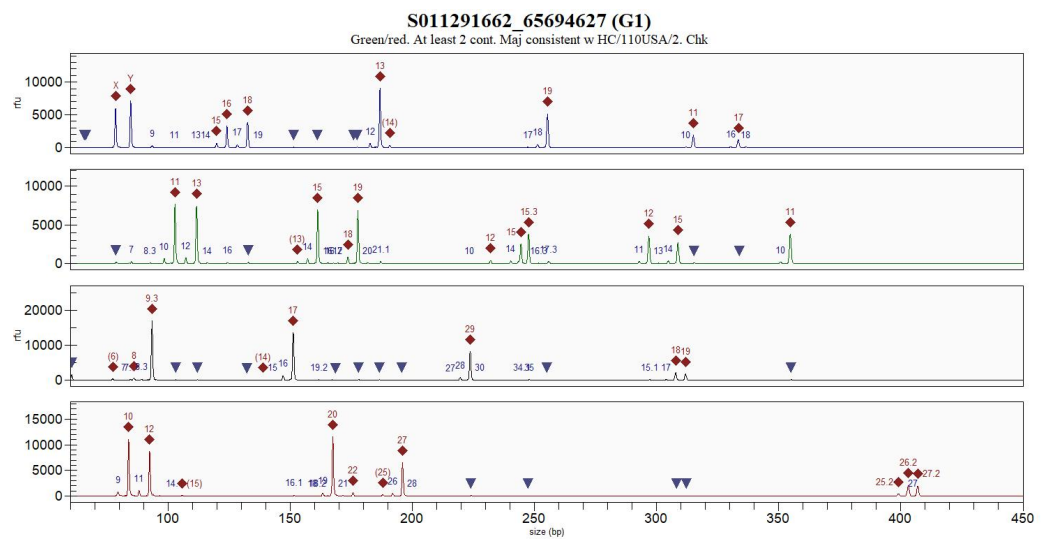

A mixed profile with an incomplete major profile matching suspect (billion times) and minor DNA components from at least two others recovered from the tool handle.

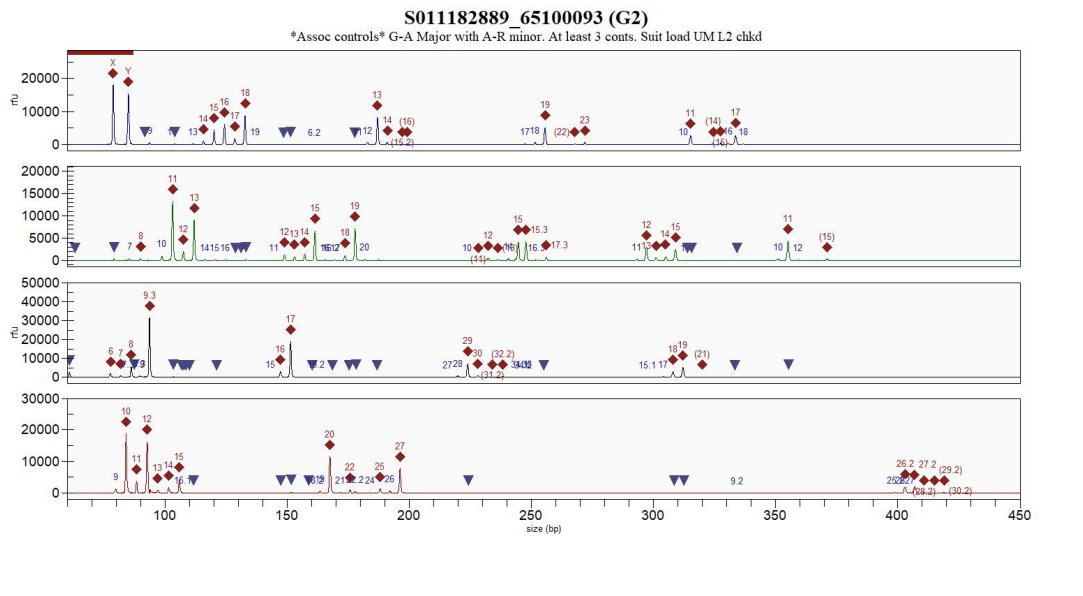

17) Assumptions

- In this case it is accepted that the mark on the window frame was made by the offender
- It is accepted that the DNA which matches the suspect has originated from him i.e. source level

18) The propositions:

- Suspect entered the property via a window taking tools which were used in the commission of the burglary and then left the tools at the property.
- Suspect was not present at the time of the incident and an offender who had previously stolen the suspects tools used them in the commission of the burglary

**Example 6**

What are the allegations made by the prosecution and the defence?

- Two suspects observed running from a carpark and arrested – both have made no comment in interview.
- Police attend and find a damaged vehicle that has had front window smashed and tools and gloves on the ground nearby

This situation is common in the UK where we are expected to comment at activity level but there is no account often until we go to court or in the days prior to the trial starting. It is often the case that we would be given a last minute account – in this case it is likely to be admit presence in carpark but deny causing damage. We would be interested to know how this would be dealt with using Bayes Nets or how much prep work would be done prior to trial.

19) What is the uncontested information?

There is no dispute that the vehicle was damaged but there is no information as to how many offenders etc

20) What is the Contested information?

Not clear, no account provided by the defence. Given that they were arrested at the scene, it would seem reasonable to assume that they would admit presence but not the offence – but where does this leave the activity of handling the screwdrivers? Could they have dropped the tools but the tools were not used in the offence? Where do the gloves fit in? Scientific support have not authorised the examination of the gloves.

The UK regulator’s recent guidance would suggest that we should assume an Hd and declare that is what we have done but this clearly risks overstating the value of the evidence and potential undermining credibility when you have to give multiple evaluations that may differ from highly significant evidence to inconclusive. The advice currently given to EFS scientists is not to make any assumptions around Hd for this reason unless it is very clear that the Hd is unlikely to change – e.g. being arrested at the scene means unlikely to deny being present.

21) Results

Both tools examined and samples taken from the handles which were subjected to DNA analysis. Both suspects could have contributed to the mixtures obtained. Evaluation using a stats package would be necessary but both suspects only have SGMPlus references to compare with DNA17 profiles.

Handle of Screwdriver 1

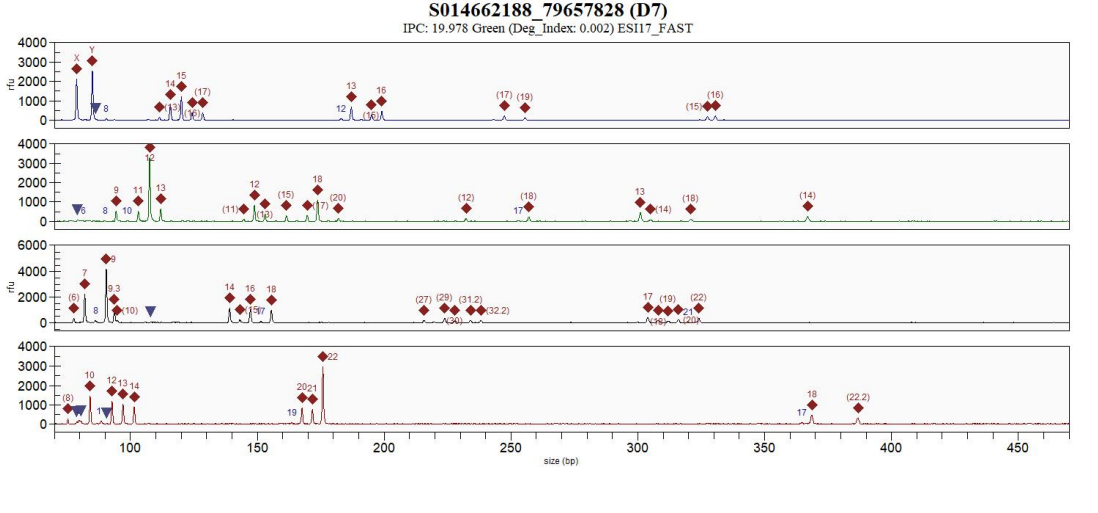

There is an additional replicate – both suspects could be contributing

Handle of Screwdriver 2

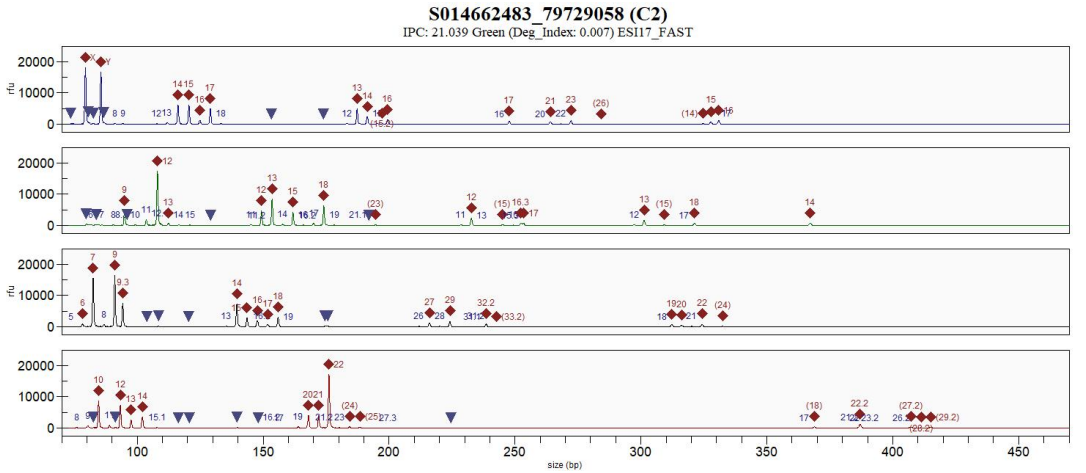

There is an additional replicate – both suspects could be contributing

Assumptions

- The assumption is that the screwdrivers were used in the offence.
- There is a possibility that one of the offenders had been wearing gloves but these have not been examined

22) The propositions:

- Hp - Suspects 1 and 2 have handled the screwdrivers believed to have been used to damage the vehicle in the carpark
- Hd – Admit being present in the carpark but deny handling the screwdrivers – the danger of proposing Hd without the detail from the suspects is that they could later admit handling but contest that the screwdrivers were used. We are also not clear whether they might admit being present when the car had been damaged or maybe they brushed against it beforehand. In reality, there are a lot of variables and the scientist needs to be very clear of the limitations

**Example 7**

23) What are the allegations made by the prosecution and the defence?

- Witness describes seeing three males steal a motorbike from a garage. The witness followed the males until the Police arrived. On police arrival the three men ran off in different directions. A short time later the suspect was arrested a short distance away. The motorbike and two crow bars were recovered from the area from which the three men ran off. The offenders were not seen to handle the crowbars but ‘they obviously took them and left them in the street’ (Direct quote from submission form). The male who was arrested denied any involvement in the burglary and then made no further comment in interview in response to the allegations against him. The suspect is a window fitter by trade.

24) What is the uncontested information?

It is accepted that there was an attempt to steal the motorbike from the garage. There is no information as to any damage to the garage and whether or not consideration has been given to tool mark comparison. The Police request is to examine the tools for handlers DNA and potentially ID another suspect.

25) What is the Contested information?

The suspect has not made any comment in relation to the motorbike or the crowbars specifically but he has denied any ‘involvement’ in the burglary.

26) Results

A mixture of DNA is obtained from the handles of both crowbars which includes DNA that could have originated from the suspect. Suspect only has an SGMPlus reference profile available. Evaluation using a statistics package would be required to obtain a statistic for source level.

Crowbar 1

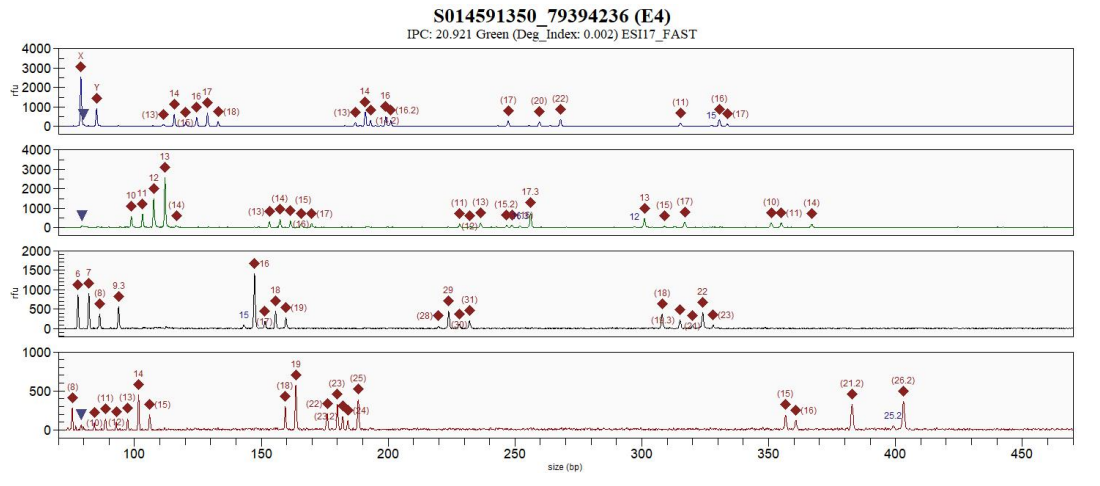

There is a replicate analysis

Crowbar 2

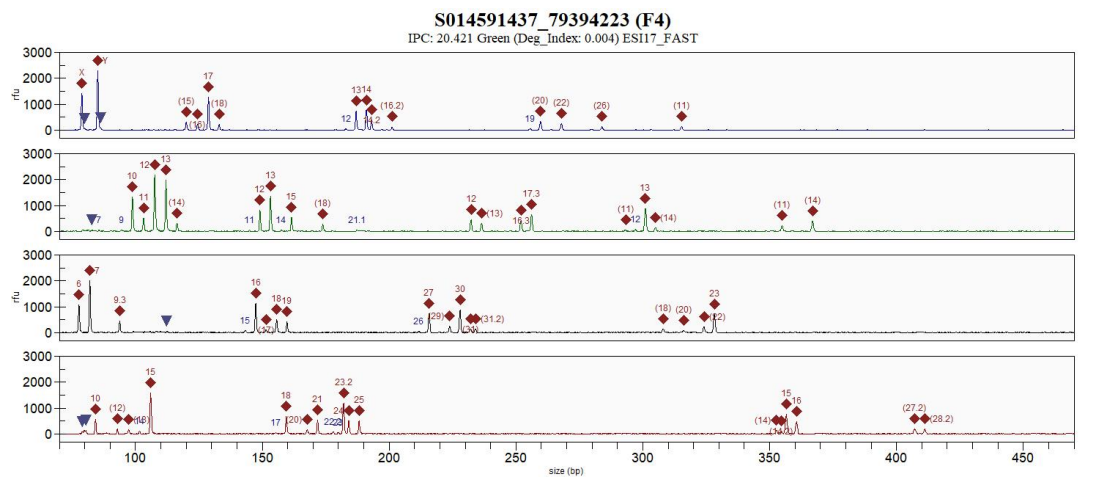

There is a replicate analysis

27) Assumptions

The assumption is that the crowbars were used in the commission of the offence. DNA is present from more than one other contributor but it is not possible to id any further offender profiles from the mixtures obtained from the crowbars.

It is an assumption that the suspect is not accepting that he had handled the crowbars. However, given his job, it is possible that he would regularly handle tools of this nature.

28) The propositions:

- Suspect handled the crowbars that are believed to have been used in the commission of the burglary.
- The suspect is not ‘involved’ – which has been taken to mean that he had not handled the crowbars.

**Example 8**

29) What are the allegations made by the prosecution and the defence?

OCG using multiple properties as cannabis grows manned by illegal immigrants who are locked into pay off their debt. Investigation identifies a hotel which when searched is found to be a large cannabis grow over multiple rooms and the electric supply had been bypassed. Two suspects are found in one of the rooms within the property but they deny involvement in the grow. A pair of pliers and multiple pairs of scissors were found at the scene along with a toothbrush, cig butt etc.

30) What is the uncontested information?

The property was being used as a cannabis grow and the suspects were present at the property

31) What is the Contested information?

The suspects were unaware of the cannabis grow and had not been involved in the cultivation or bypassing the electric supply. They have not handled the scissors or the pliers.

32) Results

A pair of pliers and a pair of scissors were examined and samples taken from the handles which were subjected to DNA analysis. Evaluation at source level requires a statistical tool for both in relation to Suspect 1.

Handle of the pair of scissors

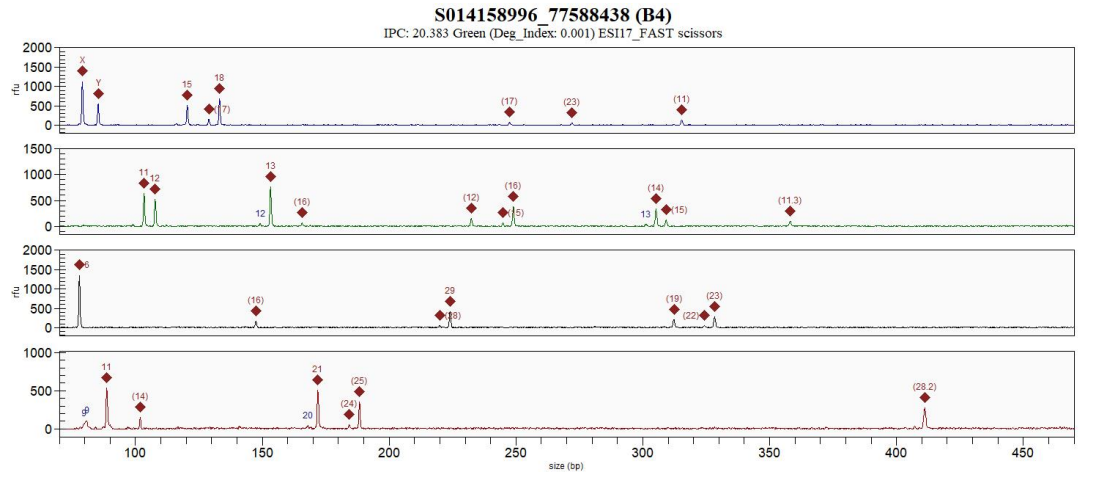

Suspect 1 is the predominating contributor, no evidence to suggest a contribution from Suspect 2. There is a replicate amp

Handle of the pliers

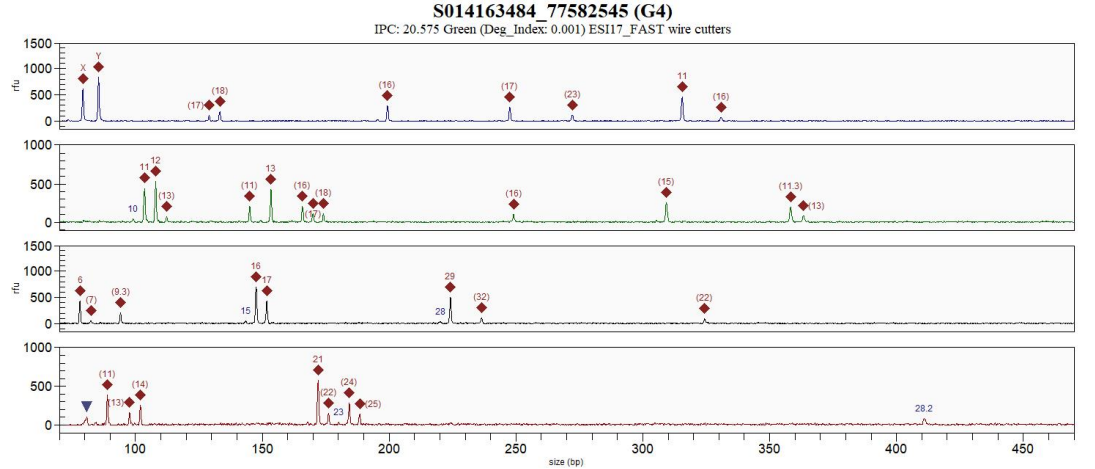

Suspect 1 is a possible contributor and nothing to suggest that Suspect 2 is contributing. There is a replicate amp

33) Assumptions

- The pliers were used to bypass the electricity and the scissors were used to harvest cannabis.
- The suspects are present in the property and therefore it would seem reasonable to assume that the DNA has originated from suspect 1

34) The propositions:

- Suspects 1 and 2 used both the plier and the scissors in the cultivation of cannabis
- Suspect 1 and 2 were staying at the property but were unaware that cannabis was being cultivated and did not use those rooms.

**Example 9**

35) What are the allegations made by the prosecution and the defence?

- Report of burglary at a school. Police arrive and a male is seen to run off.
- Offender used angle grinder to cut open a container on school property and gain access to power tools. Offender takes power tools and makes off when Police arrive. Bag with angle grinder, drill and power tools stolen from container found during area search.
- Suspect identified when seen by Police after stepping out from nearby garden. Gloves, balaclava and torch found in bushes next to the area from where the suspect became visible to Police.
- Suspect says lives locally and had been visiting a friend. He denies any part in the burglary and says that the tools, balaclava gloves and torch were not his and would not bear his DNA.

36) What is the uncontested information?

- It is accepted that a burglary took place and presumably that the angle grinder was used in the commission of the offence

37) What is the Contested information?

- Suspect denies any contact with bag or its contents

38) Results

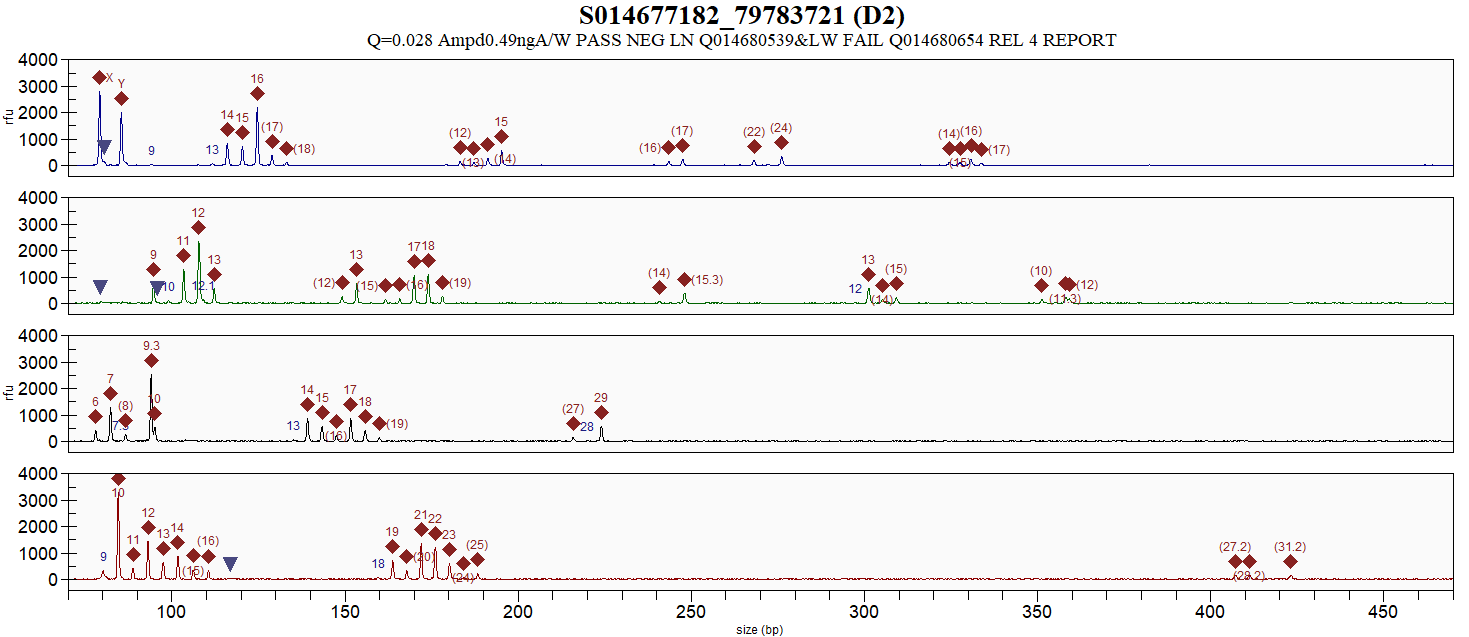

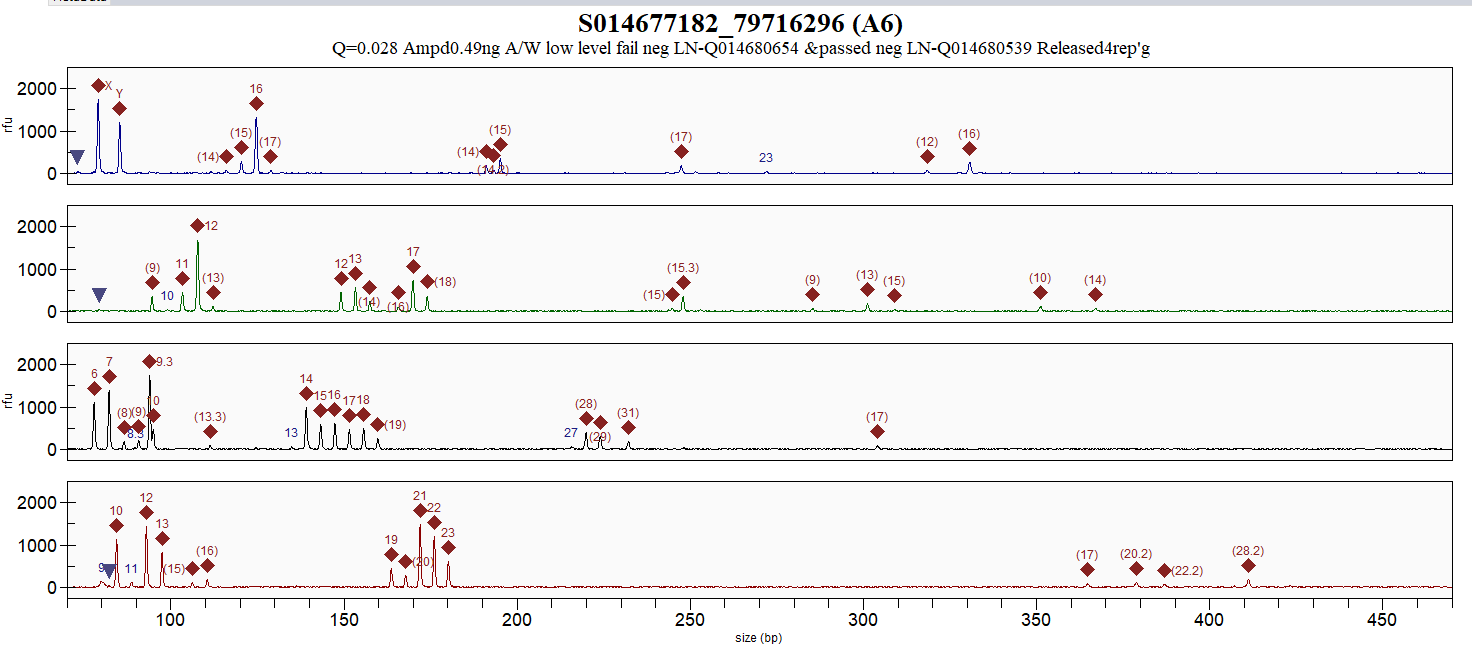
A DNA profile was recovered from the handle of the angle grinder as a mixture of multiple contributors; the suspect could be contributing. A stats package would be required to address source level.

39) Assumptions

The assumption that the angle grinder was used in the commission of the burglary.

It would not generally be the case in the UK that the source of the DNA is assumed. We cannot make assumptions that the court specifically would not accept but the Forensic Regulator would require us to comment at activity level and address handling. Therefore, EFS the statistic would be quoted to cover the source level – as specified by the Regulator and then the activity level would be dealt with separately. It must be the case that we can address activity even when source is not accepted otherwise there is a potential that evidence is massively undervalued – not so much in touch but in activity cases or bpa cases. E.g. semen on a high vaginal swab provides support of intercourse rather than social contact. This case is further complicated by the possibility that gloves were worn and the potential that the suspect would change his account.

40) The propositions:

- Suspect used the angle grinder to gain access to the container of tools
- Suspect was just in the area and has never held, touched or used the angle grinder.

Please note that results from the handle and the zip pull of the bag have been obtained. Again complex mixtures and again the suspect can be included. Whilst on paper the suspect has denied any contact, an obvious source for DNA at this level would be the bag itself and I might expect his account to change to ‘Suspect has not held the angle grinder but did see the bag and handled it’ and this is how his DNA has come to be on the angle grinder.

**Example 10**

A cannabis plantation was detected at an address 29.06.2016. Items and swabs are collected at the crime scene on the same day and sent for DNA analysis aiming to link people to this plantation. Two suspects were accused in this case, and DNA profiles matching their reference profiles were detected on several items. In court the topic of transfer and persistence were raised, and one specific hypothesis of secondary transfer was given from one of the defence lawyers.

Recovery: The items of interest were recovered at the crime scene 29.06.2016 by scene of crime officers (SOCOs).

Items received to DNA lab: 06.07.2016 (swabs collected from drinking bottles, cords, sockets, tools, fans etc. and disposable gloves and clothing).

DNA-analysis of items in questions (matching reference sample of suspect 1):

- Swab A collected from drill, stapler and pliers (all 3 items combined on one swab), found on a bench in outer room:

- DNA yield: 0.0309 ng/µL (total 6.18 ng, 200 µL extraction volume)
- DNA result: DNA-profile matching the reference samples of suspect 1

- Swab B collected from shaft of a hammer, found on bench in outer room:

- DNA yield: 0.0262 ng/µL (total 5.24 ng, 200 µL extraction volume)
- DNA result: DNA-profile matching the reference samples of suspect 1

- Swab C collected from electronic ballast and belonging cords, found on the wall in inner room:

- DNA yield: 0.0383 ng/µL (total 7.66 ng, 200 µL extraction volume)
- DNA result: DNA-profile matching the reference samples of suspect 1

- Swab D collected from electrical sockets and cords, found on the wall in inner room:

- DNA yield: 0.02 ng/µL (total 4 ng, 200 µL extraction volume)
- DNA result: DNA-profile matching the reference samples of suspect 1

- Swab E collected from electronic ballast, sockets and extension cords (combined on one swab), found in inner room:

- DNA yield: 0.037ng/µL (total 7.4 ng, 200 µL extraction volume)
- DNA result: DNA-profile matching the reference samples of suspect 1

- Trousers, found in a plastic pot in outer room. DNA sample collected from inside front pockets (sampled in the lab):

- DNA yield: 0.0149 ng/µL (total 2.98 ng, 200 µL extraction volume)
- DNA result: DNA mixture with DNA from at least three people (approx. 10:1:1 mixture), amelogenin indicated two males. The major contributor in the sample matches the reference sample of suspect 2. Suspect 1 cannot be excluded as a minor contributor.

Hypothesis, police/prosecutor: The police claim that suspect 1 deposited his DNA by direct contact.

Hypothesis, defence: The DNA from suspect 1 has been indirectly transferred from his tools (swabs A and B (hammer and drill)), maybe by someone wearing gloves, to the solid surfaces such as sockets (swabs D and E).

(This was a theory put forward by the defence in court. They did not state if the tools were stolen or borrowed, or when the transfer must have occurred. They also criticized the swabbing strategy by the SOCOs as they combined several surfaces/objects and can therefore not state from which object the detected cells were collected from)

**Example 11**

While investigating a drug case involving several addresses a weapon (rifle) was found at an address. Several swabs from different location on the weapon were collected by SOCOs and only one gave a DNA result matching the reference sample of suspect 1 (the other swabs were negative). In court the defence raised the question of indirect transfer from clothing (belonging to suspect 1) which have been wrapped around the weapon.

Recovery: The items of interest were recovered at an address crime scene 30.05.2015 by scene of crime officers (SOCOs).

Items received to DNA lab: 09.06.2015 (swabs collected from drug wraps) and 19.10.2015 (swabs collected from weapon).

DNA-analysis of item in questions (matching reference sample of suspect 1):

- Swab collected from the pipe on the rifle (comment HJ: the long cylinder part where the bullet goes through):

- DNA yield: 0.132 ng/µL (total 26.4 ng, 200 µL extraction volume)
- DNA result: DNA-profile matching the reference samples of suspect 1

Hypothesis, police/prosecutor: The police claim that the suspect deposited his DNA by direct contact while handling the weapon.

Hypothesis, defence: The DNA from suspect has been indirectly transferred from his used cloths that happened to be wrapped around the weapon.

**Example 12**

1) What are the allegations made by the prosecution and the defence?

- Mr L used a screwdriver to gain access to a vehicle from which items were stolen
- Mr L states that he has borrowed the screwdriver in the past, most recently 2 months ago, and he doesn’t know who used it to break into the vehicle but it wasn’t him.

2) What is the uncontested information?

A screwdriver was recovered from a vehicle from which items had been stolen. The screwdriver had been used to force open the door of the vehicle. The owner of the vehicle saw the screwdriver in the vehicle but did not touch it before notifying police. He did this on the day that the vehicle was broken into. Police recovered the screwdriver that same day. Mr L was questioned about this allegation 1 month after the screwdriver was recovered.

3) What is the Contested information?

Mr L says he has nothing to do with breaking into the vehicle. He has borrowed the screwdriver before from his friend to repair things at home. The last time he borrowed the screwdriver was to assemble some flatpack furniture two months ago. He hasn’t seen the screwdriver since, although he does visit his friend regularly, at least once a week, when they have beers in the garage where the screwdriver probably was.

4) Results

No body fluid testing was done

Quant trio results: SA 0.09280 ng/ul : Y 0.08750 ng/ul and approx. 0.9ng was amplified

A mixed autosomal DNA profile with a major contribution (75%) corresponding to Mr L from the handle of the screwdriver was obtained.

Remaining component, partial profile (25%) also suitable comparison should a reference become available

5) Assumptions

- DNA was present from two people
- The source of the DNA is not contested and it is agreed that the major proportion of the DNA recovered from the seized screwdriver is from L.
- The vehicle door was forced with a screwdriver by a person who was not wearing gloves
- No information on Mr L’s shedder status in known, it is considered that he or an alternative offender are compatible.

6) The propositions:

- Mr L forced open the door of the vehicle with the borrowed screwdriver
- An unknown person forced open the door of the vehicle and Mr L’s DNA was on the screwdriver from previous use.

**Example 13**

41) What are the allegations made by the prosecution and the defence?

Provide the allegations here

- Mr X participated in the throwing of a hand grenade into an apartment on June 15^th^, 2018
- Mr X states he never seen or handled a hand grenade

42) What is the uncontested information?

Describe the information that is common ground to the prosecution and the defence. Do not describe any results here

According to the information provided by the police investigator, the incident took place during the night to June 15^th^, 2018. A hand grenade was assessed to have exploded on the kitchen floor in an apartment. On the kitchen floor a safety lever to a hand grenade was found next to a stone. On the lawn outside the kitchen window a safety pin to a hand grenade was found.

43) What is the Contested information?

Describe the different positions of the prosecution and the defence. Do not describe results here.

In a hearing with the suspect Mr X on May 27^th^, 2019 he states that the last time he was inside the apartment was during a leave from prison November 2^nd^-4^th^, 2017. He was released from prison in February 2018. He also states that he has nothing to do with this and that he has never seen or handled a hand grenade.

44) Results

Results should be described here.

NFC has received LT-DNA results from a safety lever from a hand grenade. A search of the resulting DNA profile in the national DNA database resulted in a match against Mr X which was reported in a DNA database match report.

Three samples were recovered, two samples from the safety lever and one sample from the safety pin:

**The safety pin**: LT-DNA, <0,001 ng/µl, no inhibition, DNA amplification not performed

**The safety lever**: LT-DNA, 0,0015 ng/µl, no inhibition, no usable loci

**The safety lever:** LT-DNA, 0,0023 ng/µl, no inhibition, 12 usable loci. A search of the DNA profile in the DNA database resulted in a match.

The safety lever was recovered from the kitchen floor. The safety lever and the floor were covered in a dust like material which is assessed to come from a powder extinguisher that was triggered by the explosion. It is unclear how the dust may have affected the DNA result.

45) Assumptions

Based on the information, the scientist will formulate the assumptions used to evaluate the evidence.

The source of the DNA is not contested.

Mr X has not been in the apartment since November 2^nd^-4^th^, 2017.

Mr X has never handled or seen a hand grenade.

46) The propositions:

Two alternative propositions must be provided

- Mr X participated in the throwing of a hand grenade into an apartment on June 15^th^, 2018
- Mr X has never seen or handled a hand grenade

**Example 14**

1) What are the allegations made by the prosecution and the defence?

- Mr X used the screwdriver and the hammer to commit the burglary of an apartment, at a time between the 28.10.2017 and the 04.11.2017.
- Mr X states that either an unknown person wearing gloves might have stolen the tools in Poland and committed the burglary in Luxembourg or an unknown person might have stolen gloves that Mr X had used in Poland to handle the tools (that do not belong to Mr X) and commit the burglary in Luxembourg.

2) What is the uncontested information?

According to the information provided by the police investigators, the owner of the apartment was absent between the 28.10.2017 and the 04.11.2017. On the 04.11.2017 the owner discovered the burglary.

The door to the apartment was forced open. The flat is discovered in a mess. Multiple traces of searching and tool-stains (screwdriver marks, hammer marks, drill marks…) were found in the flat. The burglar had already left the apartment and there are no eye or ear witnesses. It is impossible to determine the exact time of the burglary.

Crime scene investigators discovered and swabbed multiple tools left on site by the burglar(s). The original tools were not recovered by the police investigators.

3) What is the Contested information?

According to Mr X, he has nothing to do with the burglary and he has never been in the building. During his interview on 21.02.2020, he states that he was regularly working on construction sites in Poland between 2014 and 2017. The tools are not personal. They are stored together in common storage spaces and are available for all workers.

Personal equipment, such as gloves, are stored in storage spaces accessible for all workers. As he had lost clothes, gloves or tools several times in the past, he does not pay attention if something is missing. He states that amongst the equipment, an unknown person might have either stolen his gloves or taken tools that he had used.

4) Results

Crime scene investigators swabbed multiple tools left on the crime scene - in total swabs made on 10 different tools were submitted to analysis.

6 out of 10 swabs contained no DNA. The remaining 4 swabs contained low quantities of DNA and only 2 of them provided profiles suitable for analysis.

The same DNA profile was recovered from the swabs made on one screwdriver and on one hammer. The obtained profiles were analysed as single contributor profiles that matched with Mr X's profile in a database of a neighbouring country (cf. Prüm Treaty).

5) Assumptions

- Based on the minutes of Mr X's hearing, the source of the DNA is not contested.
- Amongst other tools, a screwdriver and a hammer were used by a person to break into and to burglarize the flat.
- No specific information on Mr X's shedder status; it was considered that he or an alternative offender are compatible.

6) The propositions:

Based on the minutes of the Mr X's hearing, the judge asked us to evaluate 2 pairs of alternative propositions.

1^st^ pair of alternative propositions.

- H0 : Mr X used his screwdriver and hammer to burglarize the flat
- H1 : An unknown gloved person used Mr X's stolen screwdriver and hammer to burglarize the flat

2^nd^ pair of alternative propositions.

- H0 : Mr X used his screwdriver and hammer to burglarize the flat
- H1bis : An unknown person used Mr X's stolen gloves to handle a screwdriver and a hammer (not belonging to Mr X) to burglarize the flat

**Example 15**

1) What are the allegations made by the prosecution and the defence?

Provide the allegations here

- Mr X is in possession of three guns found in the boot of a car which he was driving
- Mr X has no knowledge of the guns, they must have been placed in the boot by someone else

2) What is the uncontested information?

Describe the information that is common ground to the prosecution and the defence. Do not describe any results here

Three firearms were found within bags (two within a black plastic bag within a cloth drawstring bag and a third inside a cloth within a white drawstring bag on its’ own) underneath a large number of coats within the boot of a car being driven by Mr X. Three men including Mr X were seen to exit the vehicle prior to the search.

3) What is the Contested information?

Describe the different positions of the prosecution and the defence. Do not describe results here.

Mr X states that he has been set up by one of the other two men that were seen to leave the vehicle. Mr X and the other man “fist bumped” each other on the morning that the guns were found and the other man regularly deposited items into the boot of the vehicle. Mr X was seen to place a bag into the boot of the car which he claims to contain his training shoes. Mr X states that there was no suggestion that he was wearing gloves and therefore, if he carried the bag containing the guns, his DNA should be on the handles.

4) Results

Results should be described here.

A full, single source DNA profile was obtained from the handgrips of one of the guns from within the plastic bag. This profile matched the reference DNA profile of Mr X.

Statistical evaluation of the match was conducted and the findings are in the region of a billion times more likely if the DNA came from Mr X than from an unknown individual, unrelated to Mr X.

(Other results from other areas of this gun and the other guns are also available)

5) Assumptions

Based on the information, the scientist will formulate the assumptions used to evaluate the evidence.

- If it is accepted that the DNA on the handgrips came from Mr X, an evaluation of how the DNA was deposited can be conducted.
- There is no additional DNA present in the result from the handgrips.
- Assumed that the correct recovery and make safe procedures have been followed.

6) The propositions:

Two alternative propositions must be provided

- Mr X handled the gun
- Mr X did not handle the gun but his DNA has been transferred to it via an unknown intermediary surface

**Example 16**

1) What are the allegations made by the prosecution and the defence?

- The accused used a screwdriver during a burglary of a flat in January.
- The defence states that the screwdriver might have been in the possession of the accused at some time, but he may have passed it on to someone else or it may have been taken by someone else. The defence also proposed that the DNA of the accused was transferred to the screwdriver indirectly that is by secondary transfer due to an unspecified contact between the accused and the real burglar.

2) What is the uncontested information?

- According to the information provided by the police investigator, the door to the flat was found open by the cleaner while the tenant was in vacation. There is no information available whether the screwdriver was used to open the door. The screwdriver was found in the bedroom. It does not belong to the tenant of the flat.

What is the contested information?

- See above: The screwdriver was used by the accused during the burglary or as the defence states, the screwdriver might have been in the possession of the accused at some time, but he may have passed it on to someone else or it may have been taken by someone else. The defence also proposed that the DNA of the accused was transferred to the screwdriver indirectly that is by secondary transfer due to an unspecified contact between the accused and the real burglar.

Results

- A DNA profile was recovered from the handle of the screwdriver that was classified as a mixed profile with a very strong major contributor. The database search with the profile of the major contributor resulted in a match with the profile of the accused. The DNA-concentration of the derived sample was appr. 180 pg/µl.

3) Assumptions

- Based on the assumption that the screwdriver belongs to the accused and based on the results described in the report from our laboratory (DNA profile of the accused is compatible with the major profile for all 16 loci), it is assumed that the source of the DNA is not contested. We have therefore considered that the major proportion of the DNA recovered from the seized screwdriver is from the accused. There is also DNA from an unknown contributor present, but only in very minor amounts (appr. 1:10).
- The screwdriver was left in the flat by the burglar. It was either used by the accused without wearing gloves or it was in his possession and use before the crime or the DNA of the accused was transferred indirectly.
- There are no specific information on the accused shedder status, it was considered that he or an alternative offender are compatible.

4) The propositions:

A1)

- The accused committed the burglary and left the screwdriver in the flat. He wore gloves during the burglary. The screwdriver was in the possession of the accused already in the time before the burglary.
- An unknown person committed the burglary and left the screwdriver in the flat. He wore gloves during the burglary. The screwdriver was in the possession of the accused at some time before the burglary, but was taken from him by the unknown person shortly before the burglary.

A2)

- The accused committed the burglary and left the screwdriver in the flat. He did not wear gloves during the burglary.
- An unknown person committed the burglary and left the screwdriver in the flat. He wore gloves during the burglary. The screwdriver was in the possession of the accused at some time before the burglary, but was taken from him by the unknown person shortly before the burglary.

B1)

- The accused committed the burglary and left the screwdriver in the flat. He wore gloves during the burglary. The screwdriver was in the possession of the accused already in the time before the burglary.
- An unknown person committed the burglary, used and left the screwdriver in the flat. The unknown person did not wear gloves. Shortly before the crime the unknown person was in contact with the accused. The way of contact between the two persons is not specified by the defence.

B2)

- The accused committed the burglary and left the screwdriver in the flat. He did not wear gloves during the burglary.
- An unknown person committed the burglary, used and left the screwdriver in the flat. The unknown person did not wear gloves. Shortly before the crime the unknown person was in contact with the accused. The way of contact between the two persons is not specified by the defence.

**Example 17**

1) What are the allegations made by the prosecution and the defence?

Provide the allegations here

- The suspect used a screwdriver (wooden handle) to manipulate stolen cars in a rented garage between the 10^th^ and the 12^th^ October
- The suspect states that he did not manipulate the cars. An unknown person used the screwdriver between the 10^th^ and the 12^th^ October

2) What is the uncontested information?

Describe the information that is common ground to the prosecution and the defence. Do not describe any results here

According to the information provided, an organized group used the garage for years. The number of members of this group is unknown. The suspect is allegedly a member of this group. The screwdriver was found in the garage.

3) What is the Contested information?

Describe the different positions of the prosecution and the defence. Do not describe results here.

The suspects states that he has nothing to do with the organized group. He never touched the screwdriver.

4) Results

Results should be described here.

A DNA profile was recovered from the screwdriver that was analysed as a mixture of two contributors consisting of major and minor profiles

5) Assumptions

Based on the information, the scientist will formulate the assumptions used to evaluate the evidence.

- Based on the assumption that the screwdriver was used by the suspect and based on the results described in the report from our laboratory (DNA profile of the suspect is compatible with the major profile for all 16 loci), it is assumed that the source of the DNA is not contested^[1]^. We have therefore considered that the major proportion of the DNA recovered from the seized screwdriver is from the suspect. There is also DNA from an unknown contributor present (minor).
- The screwdriver was used by a person who was not wearing gloves
- No specific information on the suspects shedder status, it was considered that he or an alternative offender are compatible.

6) The propositions:

Two alternative propositions must be provided

- The suspect used the screwdriver to manipulate cars between the 10^th^ and the 12^th^ October
- An unknown person used the screwdriver to manipulate cars between the 10^th^ and the 12^th^ October

**Example 18**

1) What are the allegations made by the prosecution and the defence?

- Mr S stabbed the victim
- Mr S has nothing to do with the victim’s stabbing, but he often visited victim’s home

2) What is the uncontested information?

According to the information provided by the police investigator, the door of the house wasn’t forced open. The were no witnesses of the crime. But there’s a video surveillance that shows Mr S near the property on the day of the murder. Mr S denies that he was near the property that day. The knife was recovered at the crime scene. It was identified as the murder weapon. It belongs to the victim.

3) What is the Contested information?

According to Mr S, he has nothing to do with the victim’s stabbing. During his interview, he states that he was a friend of the victim and he often visited victim’s home. They often prepared food together.

4) Results

A DNA profile was obtained from the handle of the knife. It was analysed as major/minor mixture of two contributors.

5) Assumptions

- The victim’s profile is major.
- Based on the assumption that the knife belongs to victim, presence DNA of that person wasn’t contested.
- DNA profile of Mr S is compatible with the minor profile.

6) The propositions:

- Mr S stabbed the victim
- Mr S has nothing to do with the victim’s stabbing

**Example 19**

1) What are the allegations made by the prosecution and the defence?

Provide the allegations here

- Mr X used handcrafted tools during a burglary of a restaurant on 26^th^ May
- An unknown person used the tools during the burglary of the restaurant the 26^th^ of May

2) What is the uncontested information?

Describe the information that is common ground to the prosecution and the defence. Do not describe any results here

According to the information provided by the police, a window of the building was forced open. The burglar used tape, a chain and a folding rule to craft tools in order to fish money out of a safe in the bureau. The tape used by the burglar belonged to the restaurant. Afterwards, the burglar poured orange juice over the safe and floor. The tools were recovered in the bureau by the police investigators.

3) What is the Contested information?

Describe the different positions of the prosecution and the defence. Do not describe results here.

According to Mr X, he has nothing to do with the burglary.

4) Results

Results should be described here.

A DNA profile was recovered from the chain that was analysed as a mixture of several contributors.

5) Assumptions

Based on the information, the scientist will formulate the assumptions used to evaluate the evidence.

- Based on the assumption that the chain was used by Mr. X and based on the results described in the report from our laboratory (all alleles corresponding to Mr. X. can be found in the trace (in all 16 loci)), it is assumed that the source of the DNA is not contested^[2]^. We have therefore considered that Mr. X is contributing to the trace. There is also DNA from unknown contributors present.
- No specific information on Mr X shedder status, it was considered that he or an alternative offender are compatible.

6) The propositions:

Two alternative propositions must be provided

- Mr X crafted the tools and used them to get money out of the safe on 26^th^ May
- An unknown person crafted the tools and used them to get money out of the safe on 26^th^ May

**Example 20**

1) What are the allegations made by the prosecution and the defence?

Provide the allegations here

- Prosecution: Mr X used a spoon during a burglary attempt on 7^th^ March 2018
- Defence: Mr X had nothing to do with the burglary on 7^th^ March 2018.

2) What is the uncontested information?

Describe the information that is common ground to the prosecution and the defence. Do not describe any results here

According to the information provided by the police investigator, the owner of the house returned after the burglary and found the bedroom had been ransacked. A spoon was recovered inside the wardrobe by the investigator.

3) What is the Contested information?

Describe the different positions of the prosecution and the defence. Do not describe results here.

According to Mr X, he has nothing to do with the burglary. He claimed that his DNA was somehow transferred onto the spoon.

4) Results

Results should be described here.

A DNA profile was recovered from the spoon that was analysed as a mixture of two contributors consisting of major and minor profiles

5) Assumptions

Based on the information, the scientist will formulate the assumptions used to evaluate the evidence.

- Based on the DNA profiling results generated from our laboratory (DNA profile of Mr X is compatible with the minor profile for most of the 24 loci), it is assumed that the source of the DNA is not contested^[3]^. I have therefore considered that the minor proportion of the DNA recovered from the seized spoon is from Mr X. There is also DNA from an unknown contributor present.
- The spoon was used by a person who was not wearing gloves.
- No specific information on Mr X shedder status, it was considered that he or an alternative offender are compatible.

6) The propositions:

Two alternative propositions must be provided

- Mr X used the spoon during the burglary and left it in the wardrobe on 7^th^ March.
- An unknown person used the spoon and left it in the wardrobe on 7^th^ March.

**Example 21**

1) What are the allegations made by the prosecution and the defence?

- Mr X used a stone during a burglary of a house
- Mr X states that someone using the same car as him must have transferred his DNA from the car (e.g. the steering wheel) to the stone found in the house

2) What is the uncontested information?

According to the information provided by the police, the stone was found in the house. It has presumably been used to crack the window to enter. Two young men were seen at the crime site (compatible with Mr. X who is 24 at the date of the burglary). The DNA profile from the stone matches a profile from the steering wheel of a stolen car that has been found in a parking lot of a nearby city. Presumably, this car has been used as getaway car for the burglary. The suspects' DNA profile matches both profiles, the one from the stone and the one from the steering wheel. In the weeks before the burglary, the suspect used the car almost every day, but shared it with others.

3) What is the contested information?

According to Mr X, he has nothing to do with the burglary. He states that someone else must have transferred his DNA from the car to the stone, when using the car for the burglary.

4) Results

A DNA profile was recovered from the stone that was analysed as a mixture of two contributors consisting of major and minor profiles. Peaks from the major are around 10x higher compared to the minor. A total amount of 4.5ng DNA were extracted in 75ul extraction volume.

5) Assumptions

- Based on the results from our laboratory (DNA profile of Mr. X. is compatible with the major profile for all 16 loci), it is assumed that the source of the DNA is not contested. I have therefore considered that the major proportion of the DNA recovered from the stone is from Mr. X. There is also DNA from an unknown contributor present in less quantity.
- The window was cracked by the stone.
- No information is available about whether the perpetrator was wearing gloves or not, when throwing the stone through the window.
- No specific information on Mr X shedder status.

6) The propositions:

- Mr X cracked the window of the house with the stone
- An unknown person cracked the window of the house with the stone, after getting there in the car, previously used by Mr X

**Example 22**

1) What are the allegations made by the prosecution and the defence?

Provide the allegations here

- Mr X used a cross pick during a burglary into a residence
- Mr X states that he had touched a cross pick earlier at a nearby construction site and an unknown person used it during the burglary into the residence

2) What is the uncontested information?

Describe the information that is common ground to the prosecution and the defence. Do not describe any results here

According to the information provided by the police investigator, the door to the balcony was forced open after repeated application of a 55 mm wide cross pick. The perpetrator searched through various containers and stole jewelry and other objects. The perpetrator left the cross pick on the balcony of the residence and left the location in an unknown direction.

3) What is the contested information?

Describe the different positions of the prosecution and the defence. Do not describe results here.

According to Mr X, he has nothing to do with the burglary. On his way home he once took a short-cut through a nearby construction site and had to clear some tools out of the way, among others a cross pick. This was the only place he came into contact with a cross pick. However, he could not say exactly when he was on the construction site and touched the pick.

4) Results

Results should be described here.

A DNA profile was recovered from the wooden handle of the cross pick that was interpreted as a mixture consisting of a major and a minor profile. The major DNA profile matches the DNA profile of Mr. X for the 12 evaluable and reproducible loci. The minor profile includes more than one contributor, the characteristics were at the detection limit and not reproducible, therefore the minor profile was not interpretable.

5) Assumptions

Based on the information, the scientist will formulate the assumptions used to evaluate the evidence.

We never provided activity level propositions and calculations in any of our cases, but retrospectively, we would formulate the assumptions for this case as follows:

- Based on our results (DNA profile of Mr. X is compatible with the major profile retrieved from the cross pick), it is assumed that the source of the DNA is not contested^[4]^. We have therefore considered that the major proportion of the DNA recovered from the cross pick is from Mr. X. There is also little DNA from one or more unknown contributor(s) present.
- No specific information on Mr X shedder status is available, it was considered that he or an alternative offender are compatible.

6) The propositions:

Two alternative propositions must be provided

We never provided activity level propositions and calculations in any of our cases, but retrospectively, we would formulate the activity level hypotheses for this case as follows:

- H0: Mr X used a cross pick during a burglary into a residence
- H1: Mr. X had touched a cross pick earlier at a nearby construction site. An unknown person used the cross pick for the burglary into the residence.

**Example 23**

1) What are the allegations made by the prosecution and the defence?

- Person A and Person B tried to entry the post office in the small town on the 19^th^ December at late evening. They used the crowbar trying to break the lock.
- Person A states that the bag with tool was probably stolen from his garage before the attempt of burglary in the Post office.

2) What is the uncontested information?

According to the information provided by the police investigator, the lock on the post office door was broken. A couple came across and saw two burglars, but they didn’t saw them well because it was dark. The burgles ran away and left the bag with tool at the post office door. Crowbar was beside the bag, and another 8 different items were in the bag.

3) What is the Contested information?

According to Person A, he has nothing to do with the attempt of burglary. During his interview on 28^th^ December, he states that he used to had bag with tool a long time ago, but he doesn’t use it anymore because it is too old. He also states that he didn’t remember where he had left the bag and that is it very possible that someone took the bag from his garage.

4) Results

A single DNA profile was recovered from bag handles. A DNA profile was recovered from the crowbar that was a mixture of two contributors (major and minor). The DNA profiles was recovered from another 8 items form bag, was the mixtures of two or three persons.

5) Assumptions

- The bag and the tools belong to Person A. The DNA profile obtained from bag match the DNA profile of Person A. Also, the major component of the DNA profile from the crowbar is from Person A. The minor component of that DNA profile is from unknown person.

6) The propositions:

- Person A tried to break the lock of the post office door, on the 19^th^ December.
- An unknown person tried to break the lock of the post office door on the 19^th^ December, using the person A’s stolen/lost crowbar.

**Example 24**

7) What are the allegations made by the prosecution and the defence?

Prosecution: Mr X has an illegal handgun in his possession.

Defense: Mr X has used the towel in which the handgun was wrapped when found by the police.

8) What is the uncontested information?

The handgun was found wrapped in a towel in a bag hidden on top of an elevator. Mr X lives in an apartment connected to the elevator. Mr X uses the elevator on regular basis.

9) What is the contested information?

Mr X states that he has never seen the gun before. However, he recognizes the towel, which he lost from the laundry room about a year ago.

10) Results

Material recovered from the surface of the gun gave rise to a DNA-profile representing a mixture of DNA from more than two persons. Comparison with the DNA-profile of Mr X and statistic evaluation established that the result is more than 1.000.000 times more likely if the major part of the DNA originates from Mr X than if this part originates from a random person from the population.

11) Assumptions

It is assumed that the source of the major component of the DNA recovered from the gun originates from Mr. X.

It is assumed that the towel belongs to Mr. X.

The gun was recovered from a place to which Mr. X had access.

12) The propositions:

The gun belongs to Mr. X.

The gun belongs to an unknown person who stole Mr X´s towel.

**Example 25**

13) What are the allegations made by the prosecution and the defence?

- Mr. Hospodarsky used two crowbars (N. 16 and 17) found in his backpack to force the door of newspaper kiosk during the burglary on 10^th^ May.
- Mr. Hospodarsky states that he has never used the two crowbars found in his backpack and that they belong to an unknown person.
- Mr. Vasutka used two crowbars (N. 16 and 17) found in Mr. Hospodarsky´s backpack to force the door of newspaper kiosk during the burglary on 10^th^ May.
- Mr. Vasutka states that he was using the crowbars, but has lost them by end of April.
- Mr. Hospodarsky and Mr. Vasutka committed the burglary on 10^th^ May together as a partners in crime.
- Mr. Hospodarsky states he doesn´t know Mr. Vasutka.

14) What is the uncontested information?

According to the information provided by the police investigator, the door to the newspaper kiosk was forced open. Immediately after the owner of newspaper kiosk found his kiosk burglarized, he called the Police. Mr. Hospodarsky was arrested as suspicious person close to the scene of crime, wearing the backpack with two crowbars, one clipper and some plastic bags with white substance. No gloves were founded in his backpack, neither in the scene of crime.

15) What is the Contested information?

According to Mr. Hospodarsky, he has nothing to do with burglary and he was walking close to the scene of crime incidentally. He states that he has never been using the tools carrying in his backpack and that these tools belong to an unknown person.

Investigators supposes that Mr. Hospodarsky used the tools carrying in his backpack to forced open the door of newspaper kiosk. (This was the first assumption, Mr. Vachutka as a potentional donor of biological material was found by genetic analysis).

According to prosecution, Mr. Vachutka used the crowbars to forced open the door of newspaper kiosk and Mr. Hospodarsky and Mr. Vachutka were partners in crime.

According to Mr. Vachutka, he was commonly using the crowbars for his purposes, but he noticed that the tools had gone missing few weeks before the burglary´s date and he has nothing to do with the burglary.

16) Results

DNA profiles were recovered from both crowbars (N. 16 and 17). These two DNA profiles were analysed as a mixture with one identic major contributor consisting of major and minor DNA profile. Minor DNA profile was analysed as unsuitable for interpretation.

17) Assumptions

Based on the assumption that the crowbars (N. 16 and 17) belongs to Mr. Hospodarsky and based on the results described in report KU-3576/2020 from our laboratory (DNA profile of Mr. Hospodarsky is not compatible with major profile recovered from crowbars), it´s assumed that the source of the DNA is contested. The major proportion of the DNA recovered from the both seized crowbars comes from unknown contributor, other than Mr. Hospodarsky.

The identic major DNA profile of the both seized crowbars was searched against the DNA database and the match with DNA profile of Mr. Vachutka for all 16 loci was found.

Based on the assumption that the crowbars (N. 16 and 17) belongs to Mr. Vachutka and based on the results described in report KU-3576/2020 from our laboratory (DNA profile of Mr. Vachutka, which is available in the DNA database, is compatible with major profile recovered from crowbars for all 16 loci), it´s assumed that the source of the DNA is not contested. I have therefore considered that the major proportion of the DNA recovered from the seized crowbars comes from Mr. Vachutka. There is also DNA from an unknown contributor or contributors present, but the minor profile is not suitable for interpretation.

The crowbars were used by a person who was not wearing the gloves.

Comment:

We aren´t used to operate with any kind of shedder status or shedder type of individual in our country. Every individual is automatically considered as a compatible right now. According to the information available in Police Database, Mr. Hospodarsky and Mr. Vachutka were already aligned to touch DNA samples from scene of crime, so we can consider that they are compatible.

We (Forensic experts in general) aren´t used to evaluate the evidence the way proposed in this compilation in our country, commonly there is only the statement about match or dissmatch in reports. Only in case of match or dissmatch of the DNA profiles reported as a mixture, we use the likelihood ratio calculation.

18) The propositions:

- Mr. Vachutka forced open the door of the newspaper kiosk with his crowbars on 10^th^ May.
- An unknown person forced open the door of newspaper kiosk with Mr. Vachutka´s lost crowbars by the end of April.

1. This caveat is generally applied to statements considering activity level propositions.
2. This caveat is generally applied to statements considering activity level propositions.
3. This caveat is generally applied to statements considering activity level propositions.
4. This caveat is generally applied to statements considering activity level propositions.
